# The TPLATE subunit is essential for structural assembly of the endocytic TSET complex

**DOI:** 10.1101/2020.08.13.249276

**Authors:** Klaas Yperman, Jie Wang, Dominique Eeckhout, Joanna Winkler, Lam Dai Vu, Michael Vandorpe, Peter Grones, Evelien Mylle, Michael Kraus, Romain Merceron, Jonah Nolf, Eliana Mor, Pieter De Bruyn, Remy Loris, Martin Potocký, Savvas N. Savvides, Bert De Rybel, Geert De Jaeger, Daniel Van Damme, Roman Pleskot

**Affiliations:** Ghent University, Department of Plant Biotechnology and Bioinformatics, Technologiepark 71, 9052 Ghent, Belgium; VIB Center for Plant Systems Biology, Technologiepark 71, 9052 Ghent, Belgium; Department of Biochemistry and Microbiology, Ghent University, 9052 Ghent, Belgium; VIB Center for Inflammation Research, 9052 Ghent, Belgium; Vrije Universiteit Brussel, Structural Biology Brussels, Department of Biotechnology, 1050 Brussels, Belgium; VIB-VUB Center for Structural Biology, Structural Biology Research Center, Molecular Recognition Unit, 1050 Brussels, Belgium; Institute of Experimental Botany, Academy of Sciences of the Czech Republic, Rozvojová 263, 16502 Prague 6, Czech Republic

## Abstract

All eukaryotic cells rely on endocytosis to regulate the plasma membrane proteome and lipidome. Most eukaryotic groups, with the exception of fungi and animals, have retained the evolutionary ancient TSET complex as a regulator of endocytosis. Despite the presence of similar building blocks in TSET, compared to other coatomer complexes, structural insight into this adaptor complex is lacking. Here, we elucidate the molecular architecture of the octameric plant TSET complex (TPLATE complex/TPC) using an integrative structural approach. This allowed us to describe a plant-specific connection between the TML subunit and the AtEH/Pan1 proteins and show a direct interaction between the complex and the plasma membrane without the need for any additional protein factors. Furthermore, we identify the appendage of TPLATE as crucial for complex assembly. Structural elucidation of this ancient adaptor complex vastly advances our functional as well as evolutionary insight into the process of endocytosis.

**Graphical abstract:** 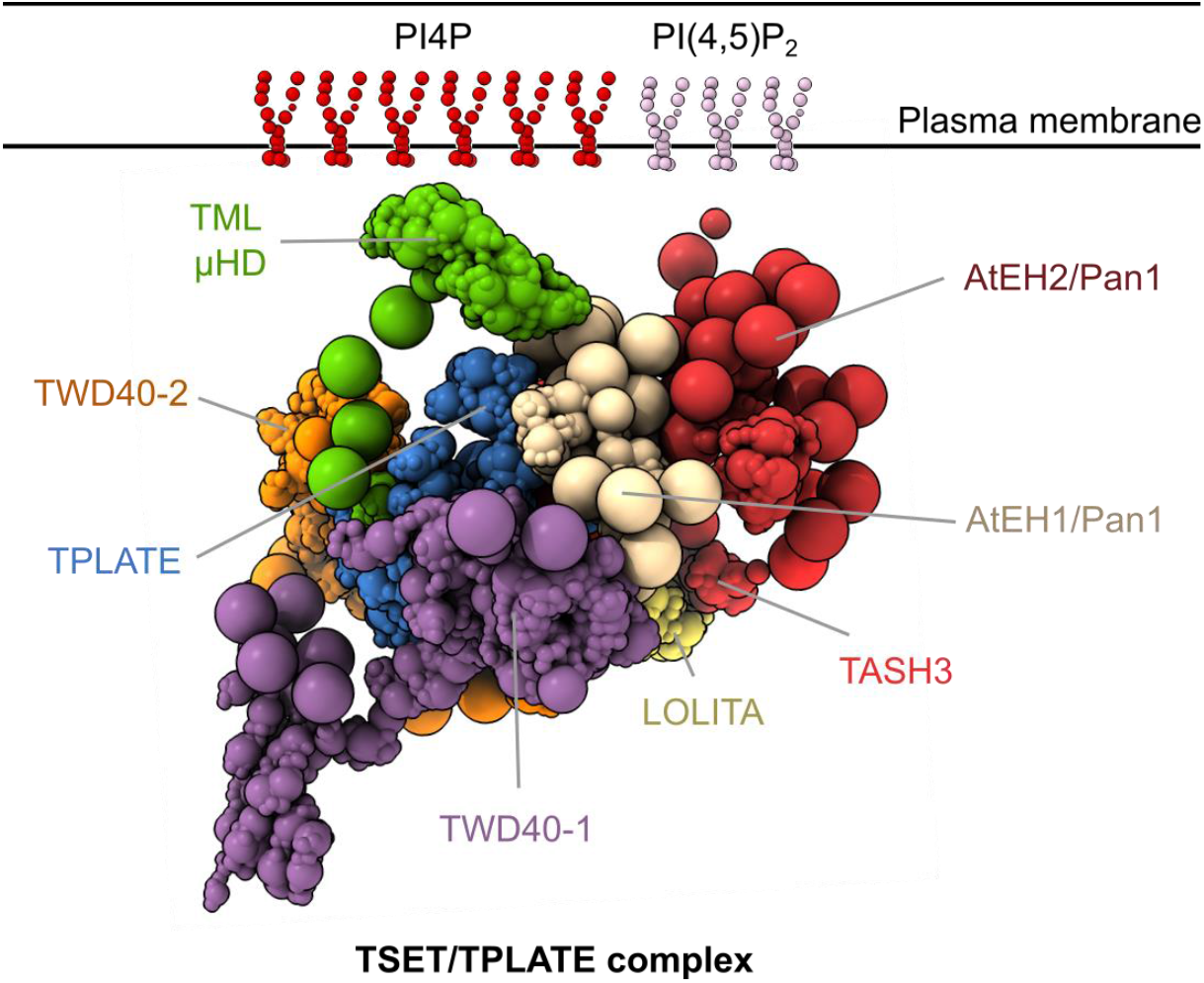

## Introduction

Eukaryotic life strongly depends on a dynamic exchange of proteins and lipids between its different organelles, a feature already present in the last eukaryotic common ancestor (LECA) (Dacks and Field, 2007). This exchange is mitigated by a wide array of protein complexes that regulate membrane shaping and coating at distinct locations. Based on the proto-coatomer hypothesis a number of complexes involved in vesicle trafficking, including the TSET complex (TSET), the Adaptin Protein complexes (AP) 1 to 5, and the Coat Protein Complex I (COPI), share a common origin (Field et al., 2011). Through time, diversification, specialization, and loss-of-function events occurred within various branches of the tree of life, but many structural leitmotifs (i.e. building blocks), as well as fundamental mechanisms, remain shared (Dacks and Field, 2007; Rout and Field, 2017). To understand vesicle trafficking and its various adaptations, mechanistic insight into multiple coating complexes across different species is vital. Structural and functional understanding of COPI and most AP complexes is available as they have been well studied in animal and yeast cells (Beacham et al., 2019; Béthune and Wieland, 2018). However, our knowledge concerning the ancient TSET complex remains very limited. TSET is broadly present among different eukaryotic supergroups but was hidden from previous studies due to its absence in the metazoa and fungi (Hirst et al., 2014). It was therefore described as a “jotnarlog” by analogy to the ancient hidden world Jotunheim in the Norse mythology (More et al., 2020).

TSET and its counterpart in plants, the TPLATE complex (TPC) are formed by TSPOON (LOLITA), TSAUCER (TASH3), TCUP (TML), TPLATE, TTRAY1 (TWD40-1), TTRAY2 (TWD40-2), and in the case of TPC supplemented by two AtEH/Pan1 proteins. TSET/TPC are stoichiometrically uniform (1:1) complexes as determined by quantitative mass spectrometry or blue native polyacrylamide gel electrophoresis (Gadeyne et al., 2014; Hirst et al., 2014). In analogy to other coatomer complexes from the AP:clathrin:COPI group; similar building blocks are present in TPC/TSET (Collins et al., 2002; Dodonova et al., 2015; Field et al., 2011; Ren et al., 2013; Rout and Field, 2017). The smallest and medium subunit, LOLITA and TML respectively, both contain a longing domain, which is in the case of TML extended with a μ-homology domain (μHD). This μHD is a unique constituent of TPC as it is absent in the TSET complex of Dictyostelium (Hirst et al., 2014). LOLITA and TML are embraced by an α-solenoid domain of one of the two large core-subunits, respectively TASH3 and TPLATE (Figure S1A). The solenoid domains are C-terminally extended by an appendage domain that contains unique features in TPC. The canonical appendage domain (platform/sandwich) is in the case of TPLATE conserved but extended by an additional anchor domain while in TASH3 it is exchanged for an SH3 domain. The core is associated with two TWD40 proteins that consist of two β-propellers followed by an α-solenoid domain, a key signature motif in the eukaryotic evolution and the emergence of proto-coatomer complexes (Field et al., 2011). The additional AtEH/Pan1 subunits in Arabidopsis TPC are the most structurally characterized members. They represent unique plant constituents uniting accessory protein interactions and membrane targeting via their EH domains while allowing dimerization through their coiled-coil regions (Sánchez-Rodríguez et al., 2018; Yperman et al., 2020). Originally TPC was described as a major adaptor module for clathrin-mediated endocytosis but recent insight also implicated the AtEH/Pan1 subunits as initiators of actin-mediated autophagy (Wang et al., 2019). As a central player in endocytosis and autophagy, TPC associates with a plethora of endocytic accessory proteins, cargo proteins as well as autophagy-related proteins (Arora et al., 2020; Gadeyne et al., 2014; Sánchez-Rodríguez et al., 2018; Yperman et al., 2020). TPLATE along with the AtEH/Pan1 subunits has been shown to play a major role in these intermolecular protein-protein interactions. Interactions among the TPC subunits remain, however, scarce. Next to its function as a protein-interaction hub, TPC localizes to cellular membranes to fulfil its role. The molecular nature of the TPC membrane interaction remains however largely unknown. Currently, only the second EH domain of the AtEH/Pan1 proteins has been shown to directly interact with anionic phospholipids, particularly PI(4,5)P2, a crucial phospholipid during endocytosis (Ischebeck et al., 2013; Yperman et al., 2020).

To unravel TPC complex function, formation and to expand on a possible direct membrane interaction, we utilized an integrative structural approach. Chemical cross-linking, X-ray crystallography, and nuclear magnetic resonance spectroscopy structures as well as comparative models were translated into three-dimensional representations and spatial restraints and combined using the integrative modeling platform (IMP) (Rout and Sali, 2019). Based on these restraints an ensemble of structures was calculated satisfying the input data. We validated the generated TPC structure by a variety of protein-protein interaction assays. Novel structural insight allowed positioning of all TPC subunits with high precision inside the complex, including the plant-specific AtEH/Pan1 subunits. We show, *in planta*, that TPLATE and its appendage domain are essential for complex formation due to its central location within the complex. Moreover, the interaction between the AtEH/Pan1 proteins and the TML μ-homology domain (μHD) provides evolutionary insight into the transition from the hexameric TSET complex to octameric TPC. Next to unraveling the TPC structure, we reveal a direct interaction with negatively charged phospholipids and orient the complex relative to the plasma membrane, providing the first mechanistic insight into its role in endocytosis.

## Results and Discussion

### The TPC structure reveals a central position of the TPLATE subunit

To gain understanding of the intersubunit arrangement of TSET as well as to generate novel insight into the functions of plant TPC, an integrative structural approach was implemented using IMP (Russel et al., 2012). IMP consists of a five-step process that starts with data gathering; combining experimental data, theoretical models, and physical principles. The second step is model representation. Input data is translated into restraints and/or representations. In the third step different models are generated and scored. This is followed by the fourth step where good-scoring models have to be clustered and evaluated based on the input data. The final step is the validation of the obtained structural model by data not used in the previous modeling steps (Rout and Sali, 2019).

To inform the relative orientation and arrangement of the TPC subunits, cross-linking mass spectrometry (XL-MS) was employed. TPC was purified, via tandem affinity purification, from Arabidopsis cell cultures expressing GS-tagged TML or AtEH1/Pan1 subunits. Both purifications yielded pure complexes and allowed the identification of individual TPC subunits via mass spectrometry and silver staining SDS-PAGE (Figure 1A and Supplementary dataset 2). After purification, on-bead cross-linking was performed with various BS_3_ concentrations followed by on-bead digest and mass spectrometry analysis (Figure 1B). BS_3_ was used as a cross-linker chemically linking protein amine moieties within a Cα spacing of approximately 25 Å. In total, twelve XL-MS datasets of two different baits at 1.2mM and 5mM BS_3_ were generated. A similar crosslinking profile was observed for all experimental conditions (Figure 1C). Finally, the datasets were merged, resulting in a final dataset of 30 inter- and 89 intra-subunit crosslinks (Figure 1C).

**Figure 1.**
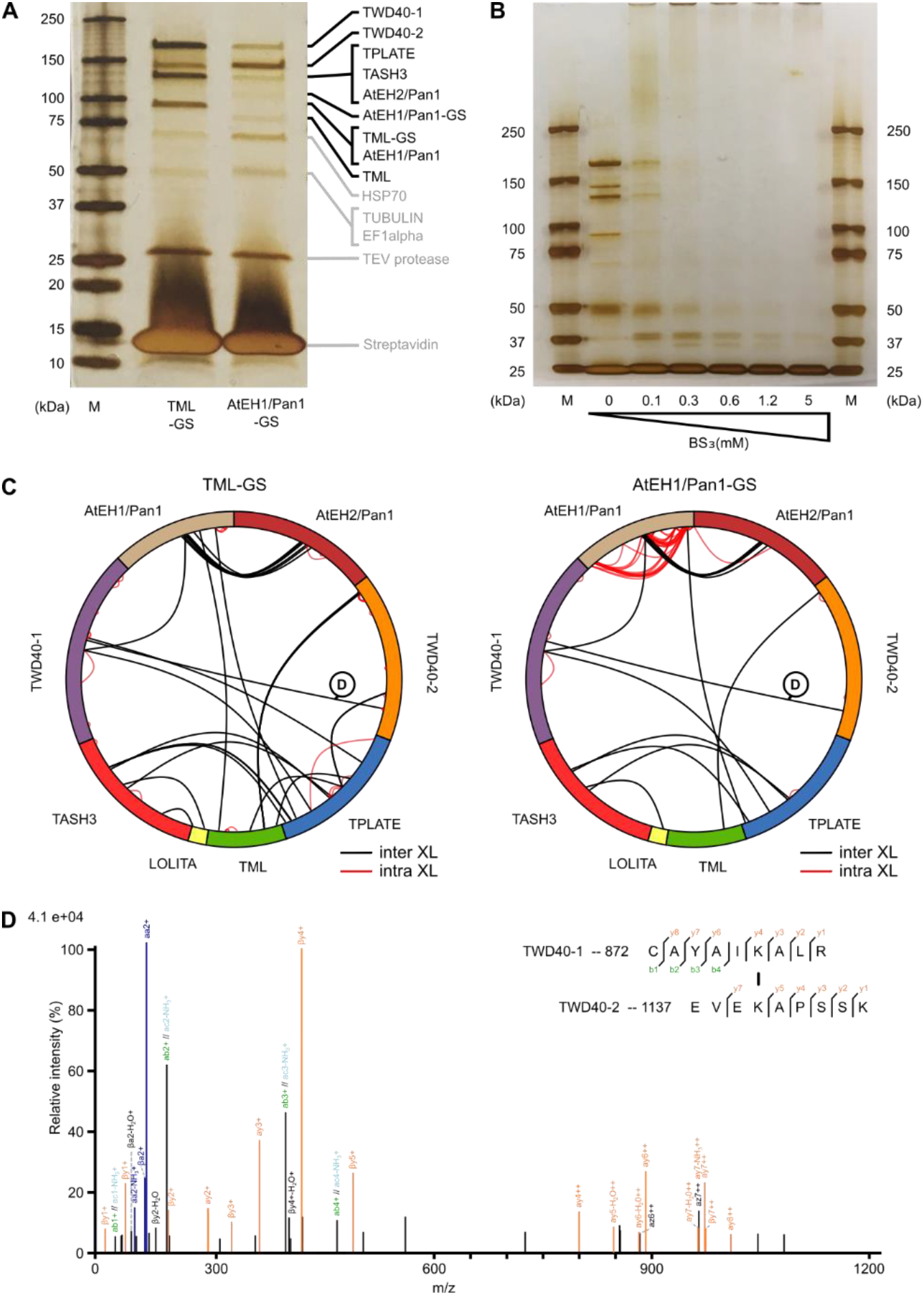
On-bead BS_3_ cross-linking reveals a highly interlinked complex. (A) Tandem affinity purification of TPC using TML and AtEH1/Pan1-tagged subunits. The purified complex was analysed by silver stain on a 4-20% SDS-PAGE. All TPC subunits, except LOLITA, can be identified based on their theoretical size and are indicated on the right side of the gel. M=Marker (B) Tandem affinity purified TML-GS before and after cross-linking with various concentrations of BS_3_, analysed by silver staining on a 4-20% SDS-PAGE gel. The vast size of TPC, with an expected molecular weight of 914 kDa, is manifested by the loss of individual subunits and the accumulation of proteins unable to penetrate the stacking gel. M=Marker (C) Cross-linking analysis following tandem purification of TML and AtEH1/Pan1 tagged subunits expressed in PSB-D cell cultures visualised by Xvis. Each analysis originates from a total of six experiments and combines 1.2 mM and 5 mM BS_3_ datasets. (D) An example of the fragmentation spectrum of the inter subunit crosslink between TWD40-1 and TWD40-2, as indicated in panel C.

Individual TPC subunits were built based on the experimental structures determined by X-ray crystallography and NMR spectroscopy or comparative models (Table S1) (Yperman et al., 2020). Given the similar domain organization in TPC, its evolutionary relationship to other coatomer complexes, and the published interaction between LOLITA-TASH3 (Figure S1), we performed protein-protein docking of LOLITA and the TASH3 trunk domain (amino acid residues 104-894) utilizing the ClusPro2.0 docking algorithm (Gadeyne et al., 2014; Kozakov et al., 2017). The best scoring model revealed an almost identical orientation of the LOLITA longin domain and the TASH3 trunk domain as in other coatomer complexes (Figure S1B). The same approach enabled positioning of the TML longin domain with respect to the trunk domain of TPLATE (amino acid residues 1-467, Figure S1B).

Input information to calculate a TPC structure included stoichiometry, chemical cross-links, protein-protein docking data as well as the experimental structures and comparative models covering 54% of the TPC sequences. Comparative models and experimentally solved structures were represented as rigid bodies while linker or indeterminate regions were described as flexible beads of different sizes ranging from 1 to 50 amino acid residues per bead (Figure S1C and Table S1). After randomization of the position of all subunits, the Metropolis Monte Carlo algorithm was employed to search for structures satisfying the input restraints. An ensemble was obtained containing 4,234 models satisfying excluded volume restraints, sequence connectivity, and at least 98% of chemical cross-links (good-scoring models). The ensemble was then analyzed and validated in a four-step protocol (Viswanath et al., 2017). Using this protocol, we tested the thoroughness of sampling, performed structural clustering of the models, and estimated the sampling precision (Figure S2A-E). All four convergence tests were passed and the analysis showed that the precision of the generated TPC structure is 39 Å as defined by the root-mean-square fluctuation of the dominant cluster containing 95% of all good-scoring models (Figure S2D). This value represents the average fluctuation of the individual protein residues or beads in three-dimensional space across the ensemble of models present in the dominant (highest-scoring) cluster.

TPC is ellipsoidal in shape with dimensions of approximately 150×160×250 Å (Figure 2A and B). The core is organized in a semi-open conformation. The trunk domains of TPLATE and TASH3 interact C-terminally and embrace the longin domains of the small and medium subunit (Figure 2B and C). This is similar to the core of the evolutionary related COPI and AP complexes (Collins et al., 2002; Dodonova et al., 2015; Ren et al., 2013). Comparable to the outer-coat complex of COPI, two TWD40 proteins overarch the core by forming a heterodimer (via their α-solenoid domains) and face with their N-terminal β-propellers the same side of the complex (Figure S2F). The TPC specific AtEH/Pan1 proteins are both attached on one side of TPC and in close proximity to the appendage domain of TPLATE, the N-terminal part of TWD40-1, and TML μHD. In line with published data, a dimerization between the coiled-coiled regions of both AtEH/Pan1 proteins was observed (Figure 2B and C) (Sánchez-Rodríguez et al., 2018; Wang et al., 2019). The localization density map for AtEH2/Pan1 and its position in the centroid structure points to a high structural flexibility of this subunit (Figure 2A and B).

**Figure 2.**
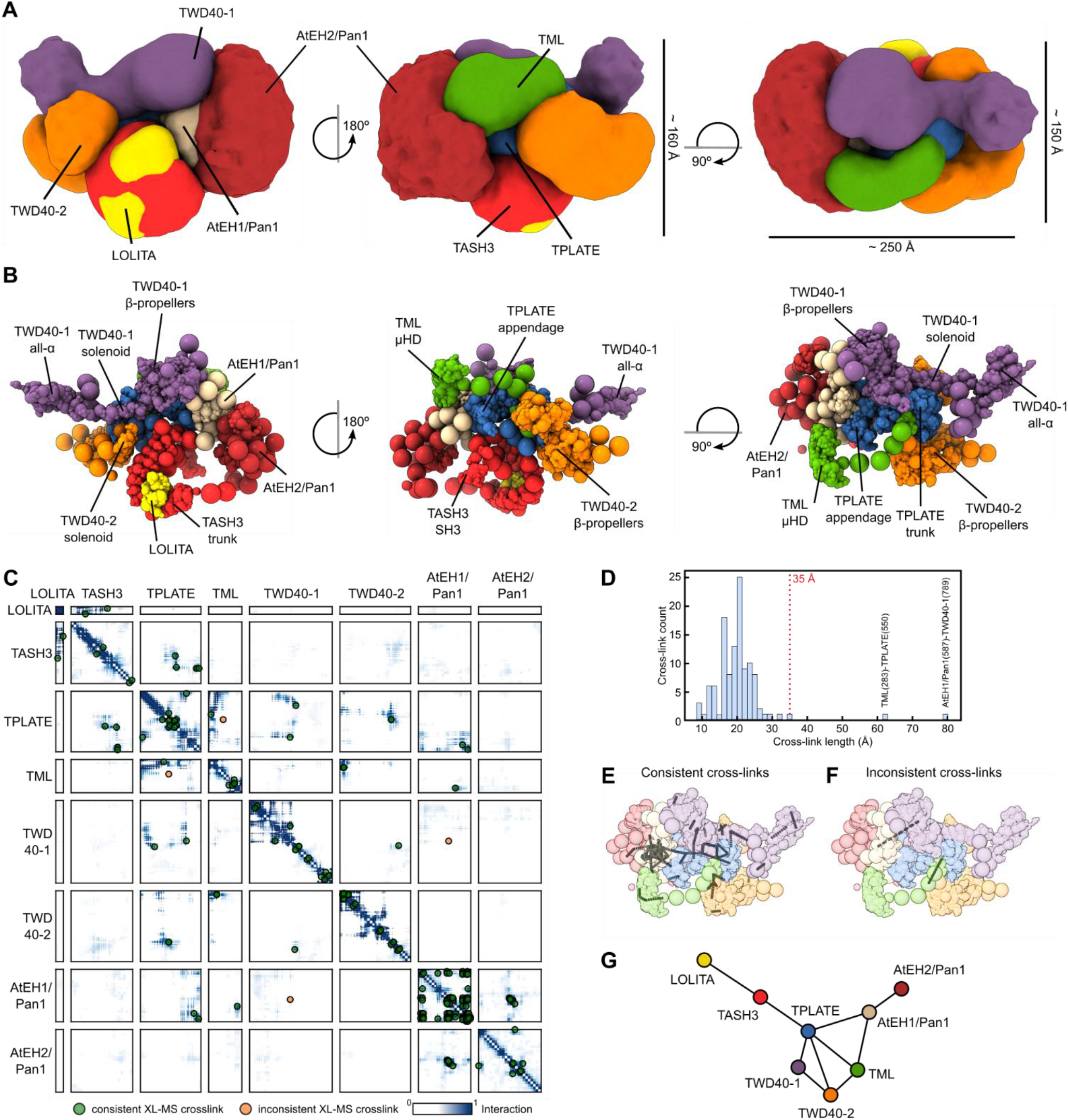
The TPC structure reveals TPLATE as highly interconnected and centrally located. (A) The architecture of TPC as obtained by the integrative modelling platform. The localization of each subunit is defined by a density map, visualized here at a threshold equal to one-tenth of the maximum. The localization density map represents the probability of any volume element being occupied by a given subunit. The approximate dimensions of TPC are 150×160×250 Å. (B) The architecture of TPC shown as a multiscale centroid structure, i.e. the structure with the minimal sum of root-mean-square deviations from all the good-scoring models in cluster 1. (C) The residue contact frequency map, calculated over 20 randomly selected models from cluster 1, is depicted by colors ranging from white (0, low frequency) to dark blue (1, high frequency). A contact between a pair of amino acid residues is defined by the distance between bead surfaces below 35 Å. Cross-links are plotted as green dots (consistent cross-links) or as orange dots (inconsistent cross-links). Each box represents the contact frequency between the corresponding pair of TPC subunits. (D) Distance distribution of obtained chemical cross-links in the centroid structure. The dashed red line represents the threshold for the consistent cross-links. Only 2 out of 129 observed cross-links are violated (located right of the 35Å border) in the TPC structure. (E) Consistent cross-links mapped on the centroid structure (grey lines). (F) Inconsistent cross-links mapped on the centroid structure (grey lines). (G) Chain-chain network diagram of the TPC structure. Nodes represent individual TPC subunits and edges (lines) are drawn between nodes which are chemically cross-linked. This reveals that TPLATE is a central hub interconnecting other TPC subunits.

Only two cross-links were inconsistent with the generated TPC structure (Figure 2C and D). These two cross-links are between AtEH1/Pan1 (K587) and TWD40-1 (K789), and between TML (K283) and TWD40-1 (789) (Figure 2D, E and F). A detailed analysis revealed that these cross-links connect disordered parts of the particular TPC subunits and the inconsistency is therefore likely caused by the coarse-grained representation of these parts limiting their flexibility.

The TPC structure together with the obtained chemical cross-links revealed the central position of TPLATE inside the complex connecting the core subunits with more auxiliary ones (Figure 2G). The TPLATE subunit can thus be seen as a hub with all its domains (trunk, appendage and anchor) forming an extensive network of interactions with other TPC subunits (Figure 2B and C).

### Yeast and *in planta* interaction data validate the obtained structure

Yeast two-hybrid (Y2H), yeast three-hybrid (Y3H) as well as *in planta* interaction methods were used to validate the obtained structure. To corroborate the protein-protein docking approach, we performed detailed Y2H mapping of the TASH3-LOLITA interaction. We confirmed that the TASH3 trunk domain is necessary and sufficient for LOLITA binding (Figure 3A). The interaction is consistent with the obtained structure of TPC. To further expand the yeast interaction landscape beyond TASH3-LOLITA, a Y2H matrix of all subunits was combined with a third non-tagged subunit, extending it to a Y3H. In contrast to previously published data, a novel weak interaction between TASH3 and TPLATE could be identified in the Y2H matrix, which was strengthened upon the additional expression of non-tagged LOLITA (Gadeyne et al., 2014). This interaction was also present between LOLITA and TPLATE, in the presence of TASH3 (Figure 3B). In accordance with the model (Figure 2G), we can conclude that LOLITA solely interacts with the trunk domain of TASH3 and that TASH3 interacts with TPLATE independently of LOLITA. Next to TASH3-TPLATE-LOLITA, a number of additional interactions were observed in Y3H. These interactions originate because the third subunit bridges two other subunits, interacting with a very low affinity or without a natural interaction surface at all. TWD40-1 interacts with TASH3 but only in the presence of TPLATE and the autoactivation of AtEH2/Pan1 is reduced in the presence of an untagged TML subunit. Both additional interactions are in line with and further validate the TPC structure (Figure 2).

**Figure 3.**
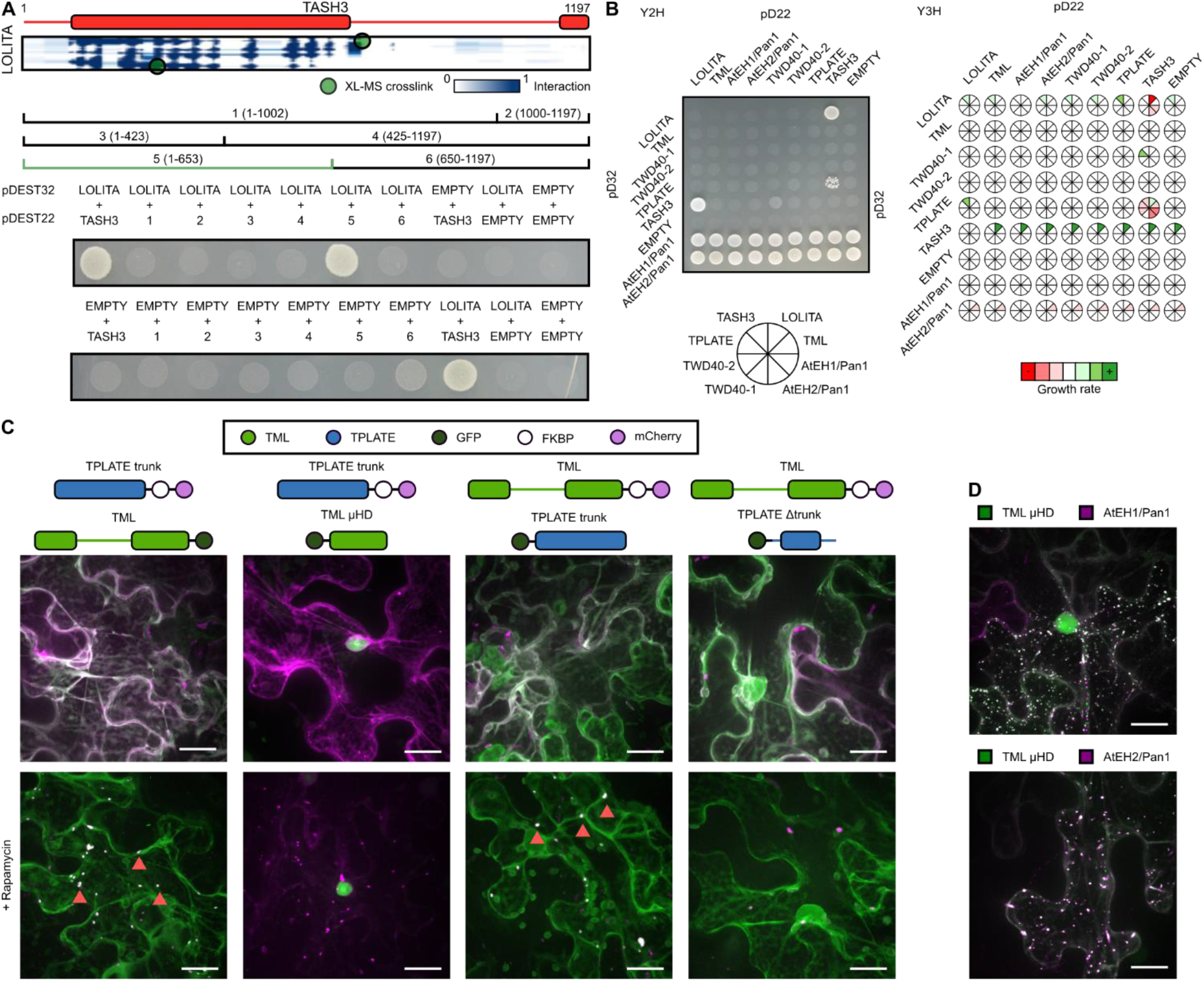
Validation of TPC structure by heterologous and *in planta* interaction assays. (A) Elucidation of the TASH3 interaction domain with the LOLITA subunit. Several TASH3 truncations (annotated as 1-6) were tested for their ability to interact with full-length LOLITA. None of the TASH3 constructs showed auto-activation when mated with an empty vector. Fragment 5, consisting of the trunk of TASH3 interacts strongly with LOLITA. The picture is representative of eight independent colonies. (B) Left - Representative Y2H matrix of all TPC subunits as well as with empty vectors used as auto-activation control. TASH3 interacts with both LOLITA as well as TPLATE. Both AtEH/Pan1 proteins strongly auto-activate. Right - Schematic visualization of the expansion of the Y2H with additional TPC subunit constructs without DNA-binding or activation-domain. Stronger interactions are indicated in green, weakened interactions as compared to Y2H are indicated in red. Full images can be found in the Supplemental Dataset 6. (C) Representative Z-stack projected images of epidermal *N. benthamiana* cells transiently expressing various GFP-fused TML and TPLATE constructs, mCherry-FKBP-fused TML and TPLATE constructs together with the MITO-TagBFP2-FRB* anchor. The used constructs are indicated above the image. Rapamycin induces re-localization of mCherry-FKBP-fused bait constructs to the mitochondrial anchor. The TPLATE trunk domain is sufficient for the TML-TPLATE interaction and TML μHD is not involved in this interaction, as it displays no co-localization. Arrows indicate co-localization of both interacting constructs at the mitochondrial anchor. (D) TML μHD is recruited to AtEHs/Pan1-induced autophagosomes. The used constructs are indicated above the image. Scale bars represent 10 μm.

To address the TML-TPLATE interaction, we utilized the recently developed knocksideway assay in plants (KSP) (Winkler et al., 2020). KSP uses the ability of rapamycin to change the localization of a bait protein and its interacting partner via hetero-dimerization of the FK506-binding protein (FKBP) and the rapamycin-binding domain of mTOR (FRB). Using KSP, it was previously shown that full-length TPLATE can re-localize together with full-length TML (Winkler et al., 2020). To this end, we transiently co-expressed various full-length and domain constructs of TML and TPLATE fused either to FKBP-mCherry or GFP, together with mitochondria-targeted FRB. We observed that the TPLATE trunk domain is sufficient for the TML-TPLATE interaction and found that the interaction is not driven by the μHD of TML (Figure 3C).

A distinctive feature of TPC compared to the TSET complex in Dictyostelium is the presence of two AtEH/Pan1 proteins. In Arabidopsis, AtEH/Pan1 proteins play a dual role, where on one hand, they drive actin-mediated autophagy, and on the other hand, they bind auxiliary endocytic adaptors as well as the plasma membrane (Wang et al., 2019; Yperman et al., 2020). Distinctive features of the AtEH/Pan1 subunits such as autophagy-interacting motifs and general domain organization are common with their evolutionary counterparts in animals (Eps15/Eps15R) and yeast (Ede1p and Pan1p). We previously showed the ability of AtEH/Pan1 proteins to recruit other TPC subunits to autophagosomes, which are formed upon overexpression of AtEH/Pan1 proteins in *N. benthamiana* (Wang et al., 2019). Here, we took advantage of this method to visualize interactions and quantitatively analyzed autophagosomal recruitment of different TPC subunits, independently for each AtEH/Pan1 subunit (Figure S3). Our quantitative analysis revealed clear differences between the recruitment of distinct TPC subunits. We found that LOLITA and TWD40-2 are the least recruited subunits by both AtEH/Pan1 proteins, in accordance with minimal contacts observed between these TPC subunits (Figure 2A). On the contrary, TWD40-1 showed the strongest autophagosomal recruitment upon overexpression of AtEH1/Pan1 consistently with the close proximity of these two subunits in the TPC structure. Finally, in the case of AtEH2/Pan1, TML and TWD40-1 were preferentially recruited, again in accordance with their position in the TPC structure. We hypothesize that the observed differences reflect on pairwise interactions between subunits of the hexameric TSET complex and the AtEH/Pan1 proteins. On one hand, the subunits, which do not exhibit any or a limited number of direct interactions are recruited to the autophagosomes only after being built in the endogenous complex. On the other hand, the subunits which can directly interact with AtEH/Pan1 proteins, are recruited to autophagosomes via their respective interacting domains, independently of complex assembly.

### The TML μ-homology domain bridges membrane and TPC subunits

One of the most significant differences between TSET in Dictyostelium and plant TPC is the presence of a C-terminal μ-homology domain (μHD) in TML, which was evolutionarily lost in the TCUP subunit of Amoebozoa. As previously hypothesized, concomitantly with this loss in Dictyostelium, the TCUP subunit lost some of its functions and the connection to other subunits (Hirst et al., 2014). A tight interaction between TML and the AtEH/Pan1 proteins is present as high temporal resolution spinning disc data shows an identical recruitment to the plasma membrane of AtEH proteins and the core subunits of TPC. This complements the data obtained in this study via Y3H and *in planta* protein-protein interaction assays that TML associates with the AtEH/Pan1 proteins (Figure 2B and Figure S3B). Previously published data hinted at a specific role of TML μHD in this interaction, as a C-terminal truncation of eighteen amino acids resulted in the loss of AtEH/Pan1 proteins, but not other TPC subunits (Figure 4A)(Gadeyne et al., 2014). To further corroborate this link, we transiently overexpressed TML μHD with either AtEH1/Pan1 or AtEH2/Pan1 in *N. benthamiana*. In line with previously shown data, the AtEH/Pan1 subunits are present on autophagosomes. The μHD did not alter the AtEH/Pan1 localisation but was recruited in both cases to the autophagosomes (Figure 3D and Figure S3). Quantification of the recruitment, by comparing the cytoplasmic signal versus colocalization with AtEH/Pan1, revealed a significant recruitment in comparison to the more distant LOLITA subunit. In conclusion, the longin domain of TML interacts with the trunk domain of TPLATE, and is coupled via a long flexible linker to its μHD that is able to associate with both AtEH/Pan1 proteins. In plants, μHD therefore acts as a bridge between the TPC hexamer and the plant specific AtEH/Pan1 subunits.

**Figure 4.**
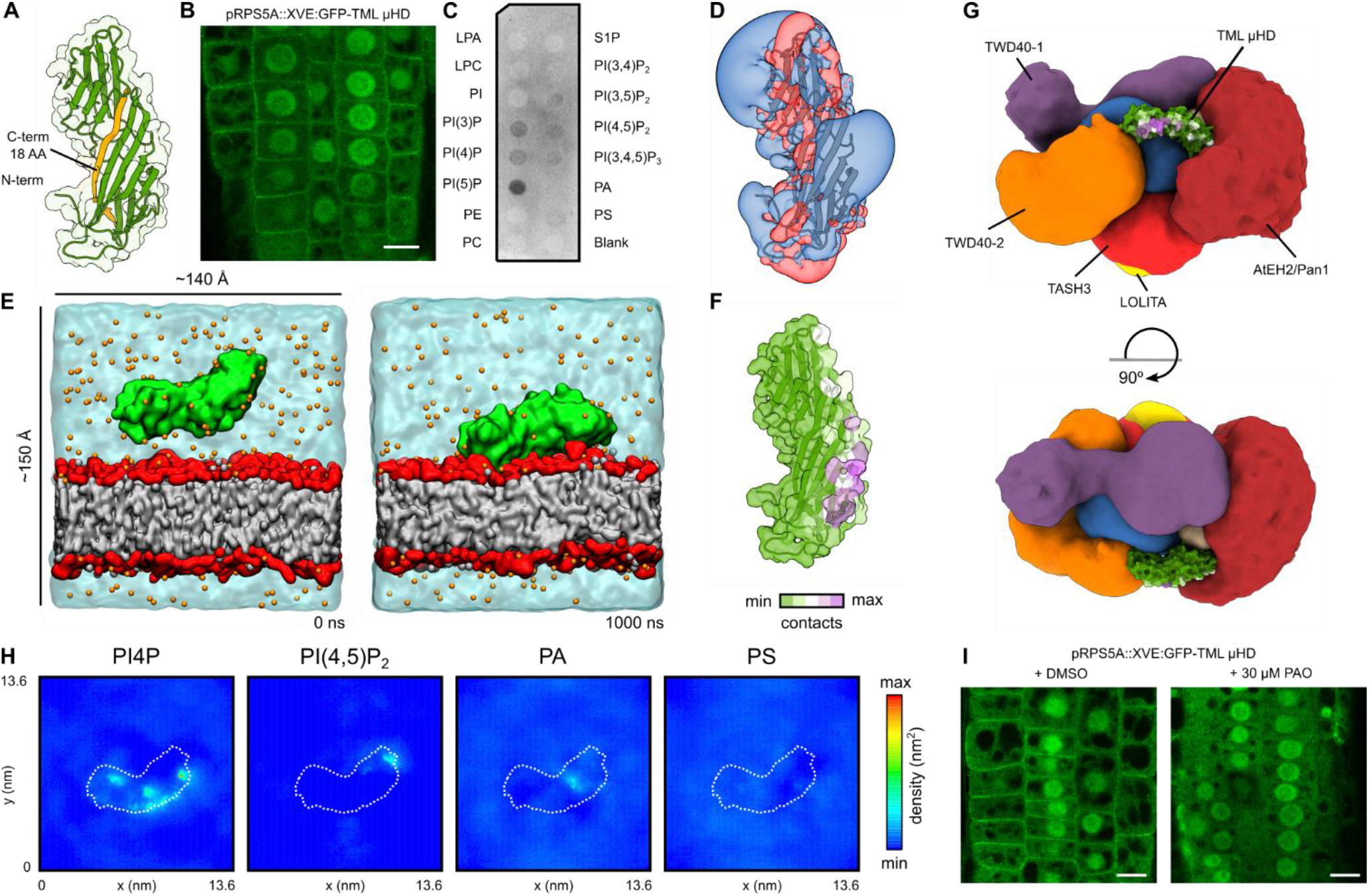
TML μHD bridges other TPC subunits with the negatively charged lipid bilayer. (A) Comparative model of the μHD of TML. The solvent-excluded surface is shown in a transparent solid representation. The c-terminal 18 amino acid (AA) truncation as published before, is shown in orange (Gadeyne et al). (B) Inducibly expressed (48 hours) TML μHD as an N-terminal GFP fusion in Arabidopsis epidermal root cells shows clear plasma membrane recruitment next to cytoplasmic and nuclear localization. The scale bar represents 10 μm. (C) PIP strip binding of TML μHD obtained by heterologous expression in *E. coli* shows clear binding to negatively charged phospholipids. Lysophosphatidic acid (LPA), lysophosphocholine (LPC), phosphatidylinositol (PI), phosphatidylinositol 3-phosphate (PI3P), phosphatidylinositol 4-phosphate (P4P), phosphatidylinositol 5-phosphate (PI5P), phosphatidylethanolamine (PE), phosphatidylcholine (PC), sphingosine 1-phosphate (S1P), phosphatidylinositol 3,4-bisphosphate (PI(3,4)P2), phosphatidylinositol 3,5-bisphosphate (PI(3,5)P2), phosphatidylinositol 4,5-bisphosphate (PI(4,5)P2), phosphatidylinositol 3,4,5-trisphosphate (PI(3,4,5)P3), phosphatidic acid (PA), phosphatidylserine (PS). (D) Electrostatic potential around the structure of TML μHD. The electrostatic potentials are represented by means of positive (transparent blue) and negative (transparent red) isosurfaces at +2 kT/e and −2 kT/e, respectively. (E) Representative snapshots of the CG-MD simulations of TML μHD with a complex negatively charged membrane. Two different timepoints are shown: the initial conditions on the left side (0 ns) and the membrane-bound protein on the right side (1000 ns). TML μHD is coloured in green, acyl chains are grey, headgroup atoms are red, sodium atoms are orange and water molecules are transparent cyan. (F) The mean number of μHD–lipid contacts of the dominant membrane orientation mapped onto the protein structure. The contacts were defined as the number of phosphate groups within 0.8 nm of protein atoms calculated over the last 500 ns. The solvent-excluded surface is shown in a transparent solid representation. (G) TML μHD superimposed onto the structure of TPC. The localization density of TML is not shown for sake of clarity. TML μHD is coloured according to the contacts with the complex membrane. (H) Two-dimensional density of different lipid molecules in the leaflet adjacent to TML μHD, calculated over the last 500 ns of the simulation, shows preferential clustering of PI4P. (I) The 30 min addition of PAO (a PI4-kinase inhibitor) resulted in complete loss of TML μHD membrane localization. Scale bars represent 10 μm.

μHDs are a common feature among vesicle trafficking complexes and are not only known to be involved in both accessory proteins and cargo interactions but have also been shown to directly interact with lipid membranes (Dergai et al., 2010; Henne et al., 2010; Jackson et al., 2012; Reider et al., 2009; Shimada et al., 2016). An unequal expression of TML and TPLATE in a double complemented, double mutant Arabidopsis line resulted in the dynamic recruitment of TML to the plasma membrane without the presence of TPLATE but not the other way around (Wang et al., 2020). Therefore, we hypothesized that TML μHD might provide simultaneous membrane recruitment and association with the AtEH/Pan1 subunits. To further elucidate its role in TPC, the μHD of TML was N-terminally fused to GFP and inducibly expressed in Arabidopsis and imaged via confocal microscopy. Next to a nuclear and cytoplasmic localization, the TML μHD was clearly recruited to the plasma membrane (Figure 4B). To rule out that the recruitment occurs through other auxiliary interactions, we analysed the protein-lipid interaction *in vitro*. We heterologously expressed and purified the domain as a N-terminal GST fusion in *E.coli*. Using a protein-lipid overlay assay, we confirmed that μHD is able to bind negatively charged lipids (Figure 4C).

Comparative modelling of the TML μHD structure revealed several positively charged patches indicating one or multiple possible binding modes towards a negatively-charged lipid bilayer (Figure 4D). To further address the TML μHD-lipid interaction, we performed extensive coarse-grained molecular dynamics (CG-MD) simulations, as this approach was shown to be a highly efficient tool to predict the membrane bound state of peripheral membrane proteins (Yamamoto et al., 2020). The simulated system contained one molecule of TML μHD, water, ion molecules and a complex lipid bilayer with the composition of charged lipids corresponding to the plant plasma membrane (Furt et al., 2010). We carried out 20 independent calculations with different starting velocities resulting in a total of 20 μs of simulation time. In all replicas, we observed that TML μHD was quickly recruited to the lipid bilayer and remained stably bound for the remaining simulation time (Figure 4E and Figure S4A). Analysis of contacts between protein residues and phosphate atoms of the lipid bilayer revealed three possible orientations of TML μHD towards the membrane with one orientation being slightly dominant (Figure S4B and Figure 4F). We then positioned TML μHD into the integrative TPC structure based on the TML localization density, the position of μHD in the centroid structure of TPC, and the observed cross-link with AtEH1/Pan1. We found that the dominant membrane-interacting mode is compatible with simultaneous membrane-binding and association of TML with other TPC subunits (Figure 4G and Figure S4C). We, therefore, hypothesize that TML μHD acts as a bridge between TPC subunits and the plant plasma membrane. This hypothesis is in agreement with the fact that in the absence of μHD, TPC is unable to be recruited to PM and the interaction with the AtEH/Pan1 proteins is lost (Gadeyne et al., 2014).

To further characterize the TML μHD interaction with the complex lipid bilayer, we monitored a 2D distribution of different lipid molecules in the lipid leaflet adjacent to the protein during our CG-MD simulations. We observed that TML μHD causes strong clustering of phosphoinositide 4-phosphate (PI4P) molecules and to a lesser degree phosphoinositide 4,5-bisphosphate (PI(4,5)P2) molecules. We did not observe significant clustering of other phospholipid molecules (Figure 4H). Given the fact that PI4P was described to control the plasma membrane identity in plant cells, we speculated that PI4P could be a main driving force for the recruitment of TML to the plasma membrane (Simon et al., 2016). To test the involvement of PI4P, the inducible fluorescently tagged μHD construct was subjected to phenyl arsine oxide (PAO) treatment which specifically affects PI4P levels in the plant plasma membrane (Simon et al., 2016). Short-term treatment (30min) at low concentrations (30 μM) resulted in a complete abrogation μHD membrane localization (Figure 5I) strongly supporting our conclusions based on the CG-MD results and the role of PI4P in the TML-membrane interaction.

**Figure 5.**
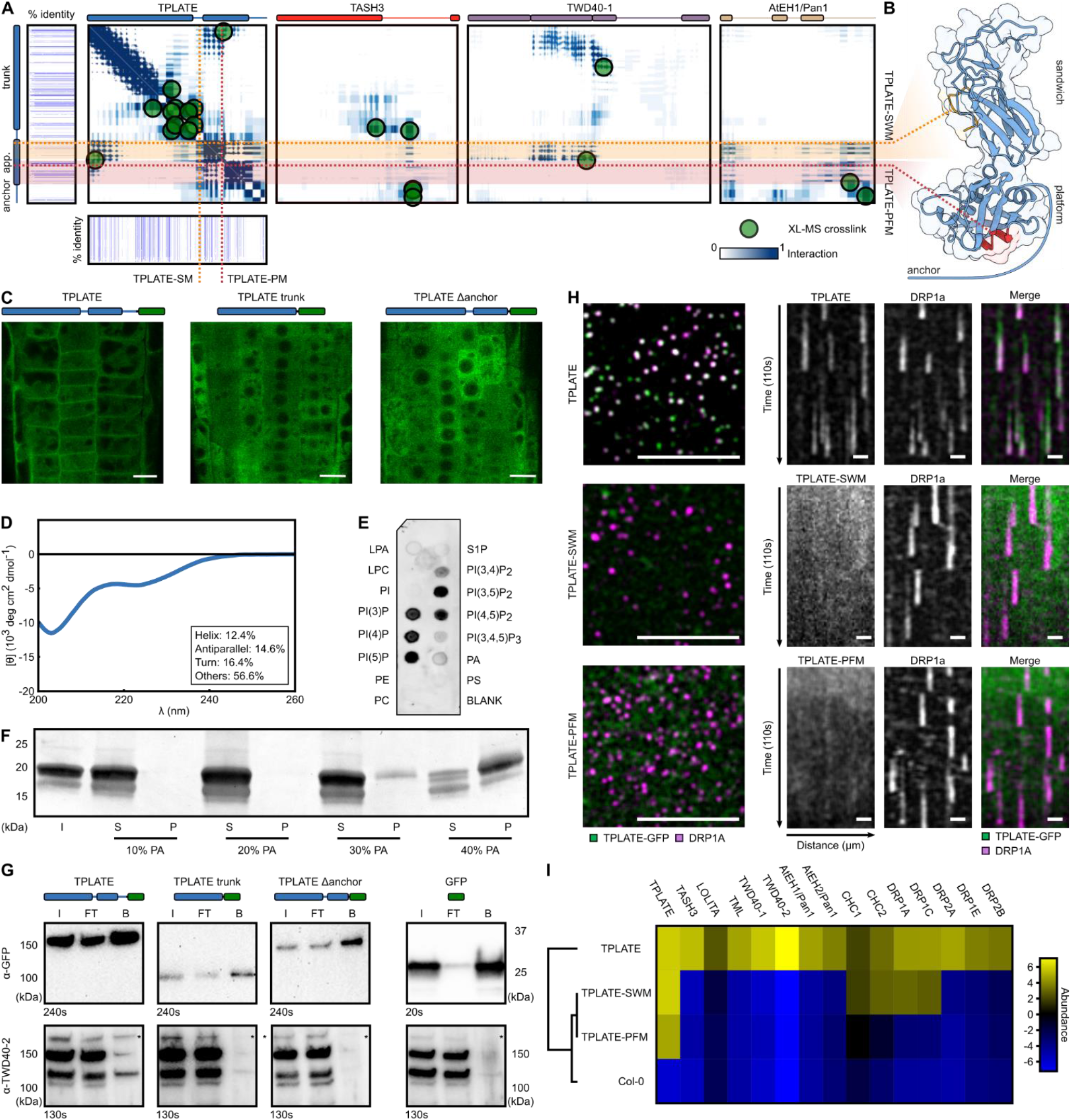
The TPLATE appendage domain is crucial for complex assembly. (A) Residue frequency contact map between TPLATE and the TASH-3, TWD40-1 and AtEH1/Pan1 subunits. The domain organisation of every subunit is shown above the contact map. The two subdomains of the TPLATE appendage, sandwich (orange) and platform (red) form an interaction hub. Cross-links are indicated as green dots. Evolutionary conservation of the TPLATE subunit is shown around the TPLATE contact map. (B) Structural features of the TPLATE appendage and anchor. Mutated residues in the sandwich (TPLATE-SWM) and platform (TPLATE-PFM) domains are indicated in orange and red, respectively. The anchor domain is shown as the light blue line. (C) *In vivo* localization of various overexpressed TPLATE truncations as analysed by confocal microscopy. The used construct is indicated above the image. Only full-length TPLATE is recruited to the plasma membrane. The absence of nuclear signal in the constructs that accumulate in the cytoplasm indicates that the truncated proteins are stable and not degraded. Scale Bar is 10 μm. (D) Circular dichroism profile of TPLATE anchor shows that the domain is mostly unstructured. The secondary structure content determined via BeStSel is shown in the inset. (E) Lipid strip of TPLATE anchor shows a binding preference for charged phospholipids. (F) Coomassie-blue stained SDS-PAGE analysis of TPLATE anchor liposome binding between various concentrations of PA (10%-40%). An increased binding capacity is observed as a higher amount of the negative charge is present. S=soluble fraction, P=pellet. (G) Co-immunoprecipitation assay comparing TPLATE with the truncated versions. Only in the case of the full-length protein an interaction with TWD40-2 (a proxy for complex assembly) could be observed. The used constructs are indicated above the image. (H) Left - TPLATE (green) and DRP1a (purple) are co-recruited at the PM as endocytic dots, whereas TPLATE-SWM and TPLATE-PFM mutants are not present in the same the focal plane as DRP1a. Scale bar is 10 μm. Right - kymographs (110s) confirm the inability of TPLATE-SWM and TPLATE-PFM to be recruited to the PM as endocytic dots. Scale bar is 10 μm. (I) Hierarchical clustered heat map of TPC subunits and identified endocytic adaptor proteins after co-immunoprecipitation with anti-GFP, as detected by mass spectrometry. Significantly different levels of proteins between TPLATE, TPLATE-SWM, TPLATE-PFM and Col-0 were determined by ANOVA followed by Tukey’s honestly significant difference test in Perseus. High and low abundant proteins are shown in yellow and blue, respectively. Mutations in the appendage domain result in the loss of TPC subunit interactions although in the case of TPLATE-SWM the interaction with some other endocytic interactors remains present. CHC stands for the clathrin heavy chain and DRP stands for the dynamin-related protein.

Although the absence of μHD still allows plasma membrane recruitment in Dictyostelium, TPC strongly depends on its presence to recruit the complex to the plasma membrane in plants. The role for PI4P specificity might play a role during certain endocytic stages as lipid conversion and the role of various phosphoinositide kinases has been implicated to regulate certain checkpoints during endocytosis in animal and yeast models (Wang et al., 2019a).

### The C-terminal domains of TPLATE are essential for complex assembly

Next to μHD, a second plant-specific modification of TSET is present at the C-terminus of the TPLATE subunit. In the plant TPLATE subunit, the appendage domain is followed by a 115 amino acid extension which we termed anchor. Our cross-linking analysis revealed that the appendage and anchor domain form a hub for intra-complex interactions (Figure 5A). Comparative modelling revealed a platform-sandwich subdomain organisation of the appendage domain while no model could be obtained for the anchor domain (Figure 5B).

To address a potential role of the anchor and appendage domain, we generated GFP-tagged TPLATE truncation constructs lacking both the appendage and anchor domain or only lacking the anchor domain and observed their localization in Arabidopsis root epidermal cells (Figure 5C and Figure S5D). Both constructs showed strictly cytoplasmic localizations in contrast to the full-length protein that was present in the cytoplasm as well as on PM and the cell plate. This is consistent with previously published data (Damme et al., 2006; Gadeyne et al., 2014). Given the loss of localization of the truncated TPLATE construct without the anchor, we hypothesized that the anchor domain could be directly involved in lipid binding. Due to the absence of a reliable homology model, we heterologously expressed the anchor domain in *E. coli*. The anchor domain eluted as a high molecular weight protein during size exclusion chromatography but size exclusion chromatography multi-angle laser light scattering confirmed its expected molecular weight (Figure S5A). This suggested that the anchor domain is loosely folded and may contain disordered regions. The circular dichroism spectrum further revealed a mostly unstructured (~70%) protein with only a very low percentage of β-sheets (~15%) and α-helices (~10%) (Figure 5D). The sequence of the anchor domain contains a highly charged region with a stretch of lysine residues. Charged unstructured regions have been implicated in membrane binding and mediating nanodomain organization (Gronnier et al., 2017). As a first proxy for membrane binding, a lipid-protein overlay assay was performed (Figure 5E). The recombinantly expressed anchor domain displayed a strong preference for charged phosphoinositides. Together with the fact that the anchor domain is highly unstructured, this preference suggested a nonspecific charge-driven interaction. To further elaborate on this possibility, liposome binding assays were performed. Comparing liposomes composed of neutral phospholipids, phosphatidylcholine (PC) and phosphatidylethanolamine (PE) to liposomes enriched with 5% PI(4,5)P2, 10% PI4P or 10% phosphatidic acid (PA), we only observed binding to liposomes loaded with 10% charged phospholipid molecules. Increasing the PI(4,5)P2 concentration to 10% also resulted in very clear binding of the anchor domain (Figure S5B and C). We next performed liposome binding assays with increasing concentrations of PA, as PA represents the simplest charged phospholipid. Higher concentrations of PA (up to 40%) indeed resulted in a stronger protein binding corroborating the non-specific charge driven interaction (Figure 5F). To test the lipid binding capacity of the anchor domain *in planta*, we expressed the domain as an N-terminal GFP fusion in Arabidopsis. However, only a cytoplasmic localization was observed (Figure S5E). Taken together with the central position of the TPLATE subunit and the fact that the anchor domain is not easily surface-accessible in the TPC structure (Figure 2A), we speculated that the anchor domain does not primarily serve as the membrane targeting module but mostly as a protein interaction hub. Consistently, we observed several crosslinks between the anchor domain and other TPC subunits (Figure 5A). To further investigate the role of the anchor domain, a co-immunoprecipitation assay was performed comparing full-length TPLATE with the truncated versions, described earlier, and probed for its ability to interact with the other TPC subunits (Figure 5G). Only in case of a full-length protein an interaction with TWD40-2 (as a proxy for complex assembly) could be observed. The TPC structure combined with the co-immunoprecipitation approach thus favours a role for the anchor domain in protein-protein interactions rather than protein-lipid interactions. Although, we cannot exclude that the anchor domain, as being intrinsically disordered, can still interact simultaneously with both lipids and other TPC subunits, especially when extended. However, such flexibility of the anchor domain is limited in the integrative TPC structure because of the coarse-grained representation of unstructured parts.

Next to the anchor domain, the TPC structure revealed a vast number of contacts between the appendage domain of TPLATE and both TWD40-1 and AtEH1/Pan1 (Figure 5A). A detailed structural analysis of the TPLATE appendage revealed a similar bilobal organisation as known appendage domains in other coatomer complexes, consisting of sandwich and platform subdomains (Figure 5B). Appendage domains were initially appointed a crucial role as auxiliary protein interaction platforms. Recent evidence based on electron microscopy in both AP-2 and COPI hints also at a role in coat formation due to the close proximity between the appendage domain and the N-terminal β-propeller of clathrin or α-COP (Dodonova et al., 2017; Kovtun et al., 2020; Paraan et al., 2020). To assess the role of the appendage domain in TPC formation, mutations were made in the evolutionary most conserved stretches of the platform (orange) and sandwich (red) subdomains (Figure 5A and B), respectively named TPLATE-PFM (PlatForm Mutant) and TPLATE-SWM (SandWich Mutant). To obtain complemented lines the GFP-tagged mutation constructs were transformed into heterozygous TPLATE T-DNA insertion lines. Expression of all constructs was validated by western blot (Figure S5F). After extensive screening no homozygous insertion line could be identified. Segregation analysis of heterozygous insertion lines revealed that both appendage mutants were unable to complement the TPLATE mutation, confirming the requirement of the appendage domain for TPLATE to function (Table 1). In addition, no membrane localization of both TPLATE SWM and -PFM constructs was observed when combined with the styryl dye FM4-64 (Figure S5G). To compare the membrane recruitment, both lines were crossed with the dynamin-related protein 1A (DRP1A) endocytic marker. Spinning-disk confocal microscopy revealed dynamic endocytic spots containing DRP1A in all TPLATE lines. Full-length TPLATE localized in the same focal plane of the DRP1A endocytic foci, indicating membrane recruitment. This was however not the case for TPLATE-SWM and -PFM mutants (Figure 5H). Given the central position of the TPLATE subunit in the TPC structure (Figure 2G), we compared the interactome of TPLATE with TPLATE SWM/PFM-mutations. Co-immunoprecipitation combined with mass-spectrometry analysis revealed the inability of these mutated TPLATE isoforms to interact with any other TPC complex subunits (Figure 6I and Figure 5SH). As revealed by XL-MS and the integrative TPC structure, the appendage domain of TPLATE is in close contact with its trunk domain to position β-propellers of TWD40-1 close to the hetero-tetrameric core (Figure 2B and C). β-propellers are known for their ability to interact with both auxiliary proteins as well as the plasma membrane. We hypothesize that the correct orientation of the appendage by interacting with the trunk domain and the TWD40 proteins is crucial for proper complex assembly and function of the TPLATE complex at the plasma membrane.

**Table 1 -.** Functionality analysis of TPLATE appendage mutants. Segregation ratios of the progeny of *tplate* heterozygous mutants expressing TPLATE-SWM and TPLATE-PFM appendage mutation constructs driven from the functional pLAT52 promotor, as analyzed by PCR analysis. In all T-DNA insertion offspring lines a 1:1 ratio of T-DNA vs WT was observed in both constructs due to male sterility. Three individual transgenic lines were analyzed for each construct. The χ2-test was used to test whether the segregation ratio deviated from 1:1. χ2 0.05 (1) = 3.841.

| Mutant lines | T-DNA | WT | Total | $\chi^2$ |
| --- | --- | --- | --- | --- |
| TPLATE-SWM-1 | 13 | 11 | 24 | 0.167 |
| TPLATE-SWM-2 | 12 | 12 | 24 | 0.000 |
| TPLATE-SWM-3 | 13 | 11 | 24 | 0.167 |
| Total | 38 | 34 | 72 | 0.174 |
| TPLATE-PFM-1 | 11 | 13 | 24 | 0.167 |
| TPLATE-PFM-2 | 8 | 14 | 22 | 1.637 |
| TPLATE-PFM-3 | 12 | 12 | 24 | 0.000 |
| Total | 31 | 39 | 70 | 0.914 |

In conclusion, we implemented a highly multidisciplinary approach to structurally characterize the evolutionary ancient TSET/TPLATE complex. By combining diverse experimental methods together with the integrative modelling platform, we demonstrate that the TPLATE subunit forms a central hub in TSET/TPC creating a vast array of protein-protein interactions and thus being indispensable for TSET/TPC assembly. The appendage domain and the plant specific anchor domain play herein a vital role. We could also link the plant-specific features of TPC. Namely, the AtEH/Pan1 proteins and the μHD of TML. Furthermore, our data clearly points to a direct interaction between the complex and the plasma membrane without the need of any additional protein factors. The generated TPC structure suggests many structural similarities between TSET/TPC and other coatomer complexes like COPI and AP2-clathrin. It will be of interest to further investigate if TSET/TPC can form higher ordered structures when bound to a membrane (i.e. a coat) similarly to other coatomers. As the integrative modelling approach is inherently an iterative process, the obtained TPC structure can be further improved as new experimental data becomes available.

---

**Figure S1.**
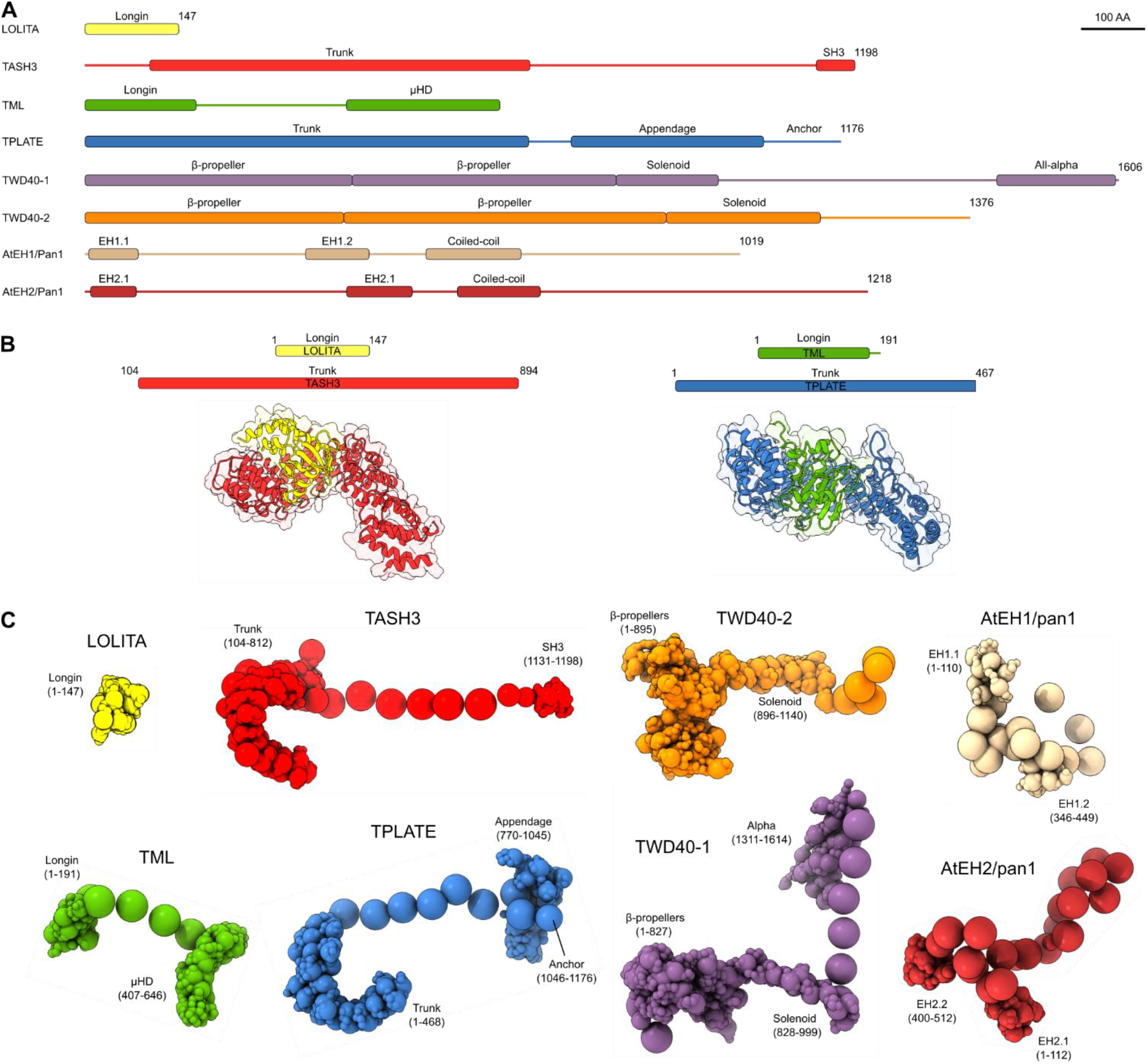
TPC domain architecture. (A) Domain organisation of the different TPC subunits based on the experimentally solved structures, comparative models and secondary structure predictions. Domains are indicated by boxes and unstructured parts of individual subunits are indicated by a solid line. The scale bar represents 100 amino acids (B) Protein-protein docking allowed to position the longin domains of LOLITA and TML into the α-solenoid domains of TASH3 and TPLATE, respectively. The amino acids used are indicated at the top of the panel. (C) Three dimensional multiscale structural representation of the various TPC subunits. The individual domains were represented by beads of varying sizes (1 to 50 amino acid residues per bead), arranged into either a rigid body or a flexible string of beads.

**Figure S2.**
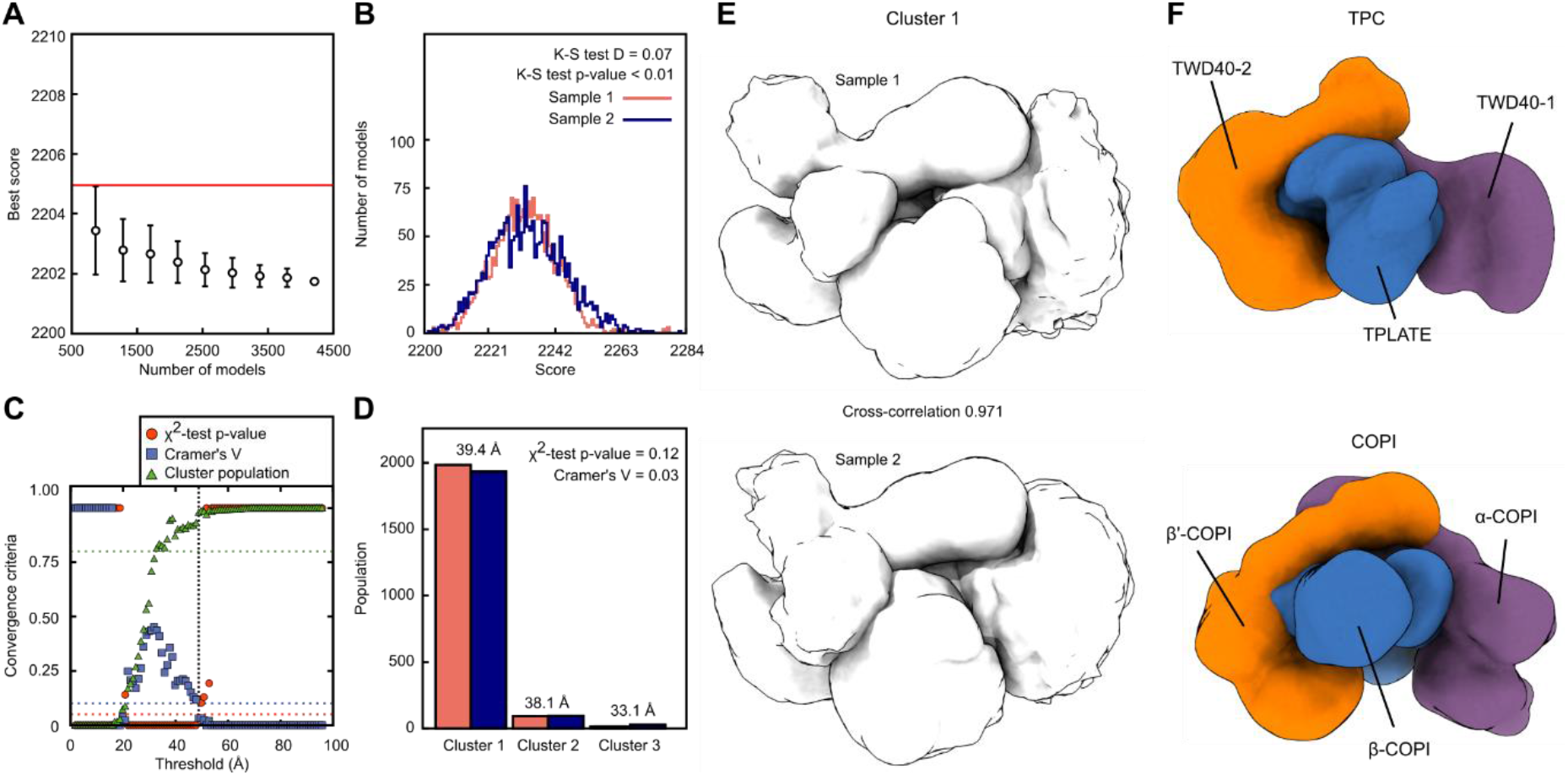
Convergence, sampling exhaustiveness and ensemble precision of the integrative TPC structure. (A) Convergence of the model score calculated for 4,234 good-scoring models. Scoring did not improve after the addition of more independent models. The error bars represent the standard deviation of the best scores, estimated by repeating the sampling 10 times. The red line depicts a lower bound on the total score. (B) Splitting of the obtained good-scoring models into two sample populations (blue and light red) resulted in significantly different score distributions (p-value < 0.01), but the magnitude of the difference is however negligible (0.07), therefore both score distributions are effectively equal. (C) The sampling precision as defined by three criteria; first, the p-value calculated using the χ2-test for homogeneity of proportions (red dots); second, an effect size for the χ2-test is quantified by the Cramer’s V value (blue squares); third, sufficiently large clusters (containing at least 10 models) visualized as green triangles. The vertical dotted grey line indicates the root mean square displacement (RMSD) clustering threshold at which three criteria are satisfied (p-value > 0.05, Cramer’s V < 0.10, and the population of clustered models > 0.80). The sampling precision is thus 49 Å. (D) Using the sampling precision as the threshold, populations of sample 1 (light red) and 2 (blue) form three clusters. 95% of the models belong to cluster 1, which has a precision of 39 Å. (E) Localization density maps for Sample 1 and Sample 2 of Cluster 1, visualized here at a threshold equal to one-tenth the maximum. The cross-correlation of the localization density maps of the two samples is 0.971, indicating that the position of TPC subunits in the two samples is effectively identical at the model precision of 39 Å. (F) A similar orientation of the two largest subunits is observed in TPC and COPI. For TPC, localization density maps of the depicted subunits visualized at a threshold equal to one-fifth the maximum are shown. For COPI (pdb code 5A1U, chain C, D and G), the molecular surface of each subunit is approximated by Gausssian surface at a resolution of 25 Å.

**Figure S3.**
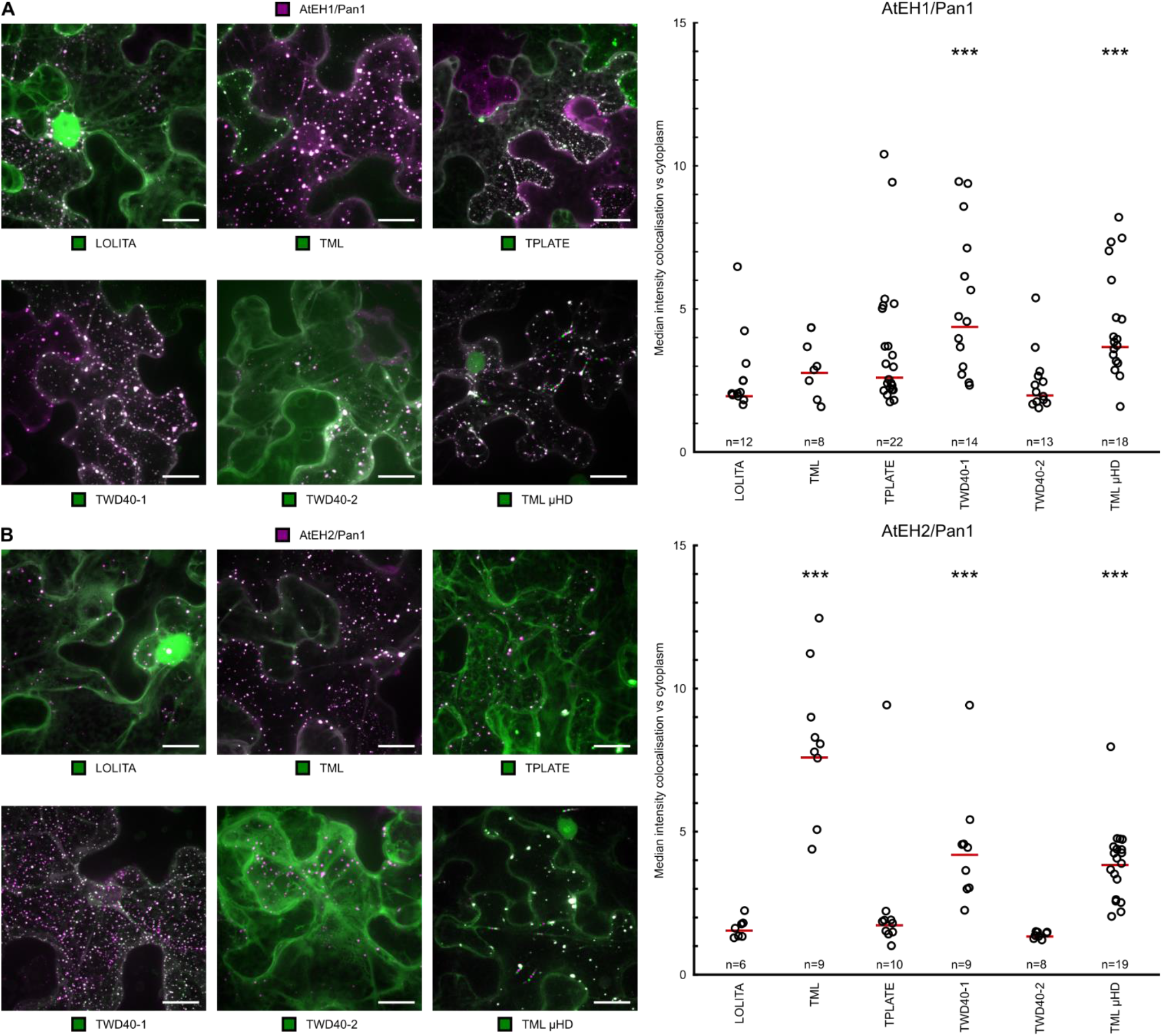
Validation of TPC structure in *N. Benthamiana*. (A-B) Representative Z-stack projected images of epidermal *N. benthamiana* cells transiently expressing different TPC subunits or TML μHD together with either AtEH1/Pan1 or AtEH2/Pan1. Quantification of TPC subunit recruitment to autophagosomes initiated by overexpression of AtEH/Pan1 subunits is shown on the right side. Recruitment is defined by the ratio between the median signal intensity in autophagosomes compared to the cytoplasm. Three stars indicate a significant difference (p < 0.05) between a given TPC subunit/domain and both LOLITA and TWD40-2 evaluated by the Tukey multiple pairwise-comparison. The red line represents the median and n depicts a number of analysed cells. Scale bars represent 10 μm.

**Figure S4.**
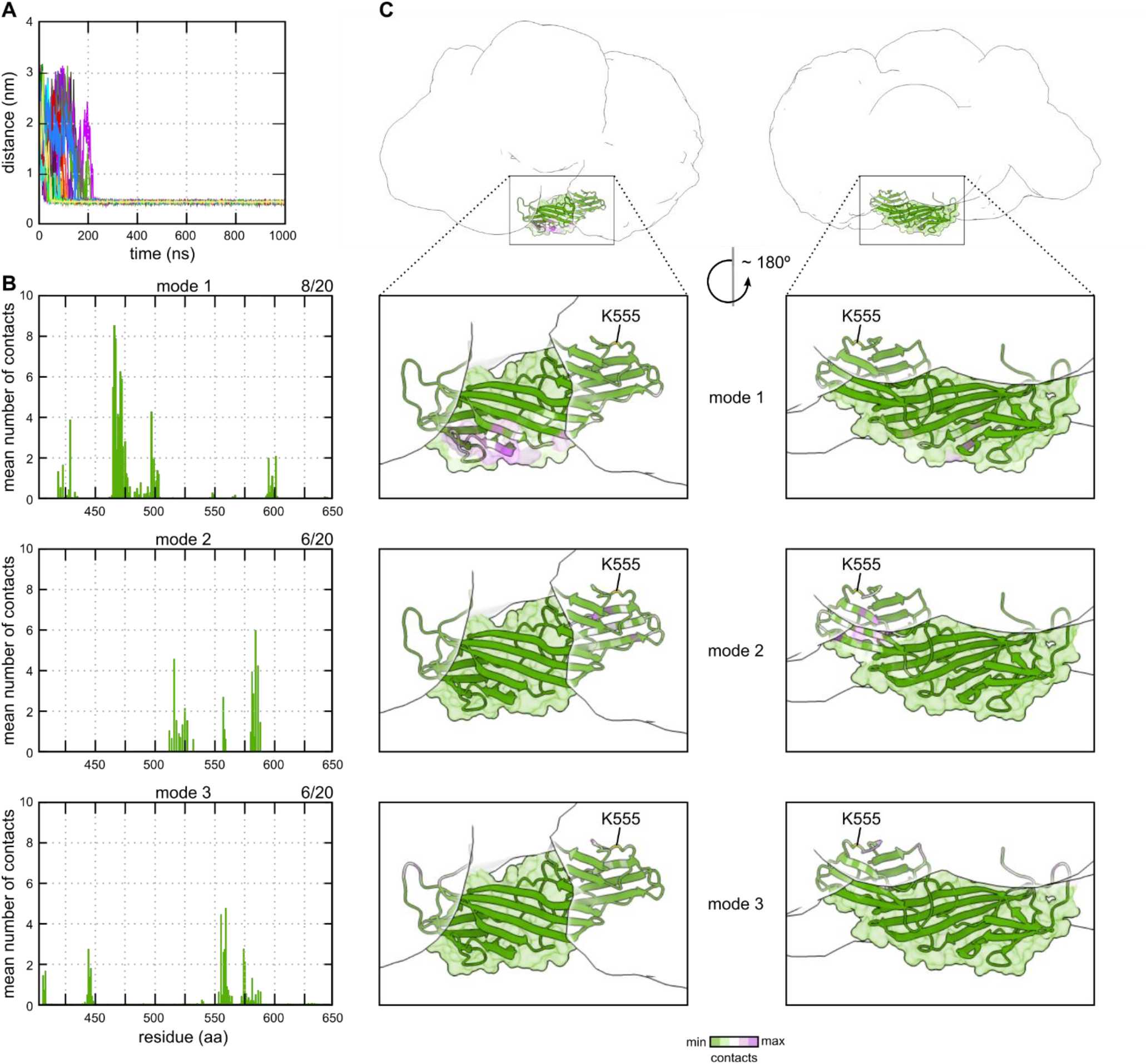
Only one lipid-binding mode of TML μHD is compatible with the simultaneous lipid and TPC interaction. (A) Progress of CG-MD simulations shown as the minimal distance between the center of mass (COM) of TML μHD and COM of the lipid bilayer. Different replicas are depicted in different colours (B) The mean number of TML μHD contacts with the lipid bilayer per residue for three identified binding modes. The contacts were defined as the number of phosphate groups within 0.8 nm of protein atoms calculated over the last 500 ns. (C) TML μHD superimposed onto the structure of TPC. Localization densities of the TPC subunits are shown in transparent white. The localization density of TML is not shown for sake of clarity. TML μHD is coloured according to the contacts with the complex membrane for the respective binding mode. TML lysine 555, identified as cross-linked to AtEH1/Pan1, is highlighted. Only binding mode 1 is consistent with the simultaneous interaction of the μHD with the lipid bilayer and with other subunits of the TPC structure. TML μHD was manually positioned into the TPC structure based on the TML localization density, position of μHD in the centroid structure of the TPC model and the chemical cross-link with AtEH1/Pan1 using the Chimera program.

**Figure S5.**
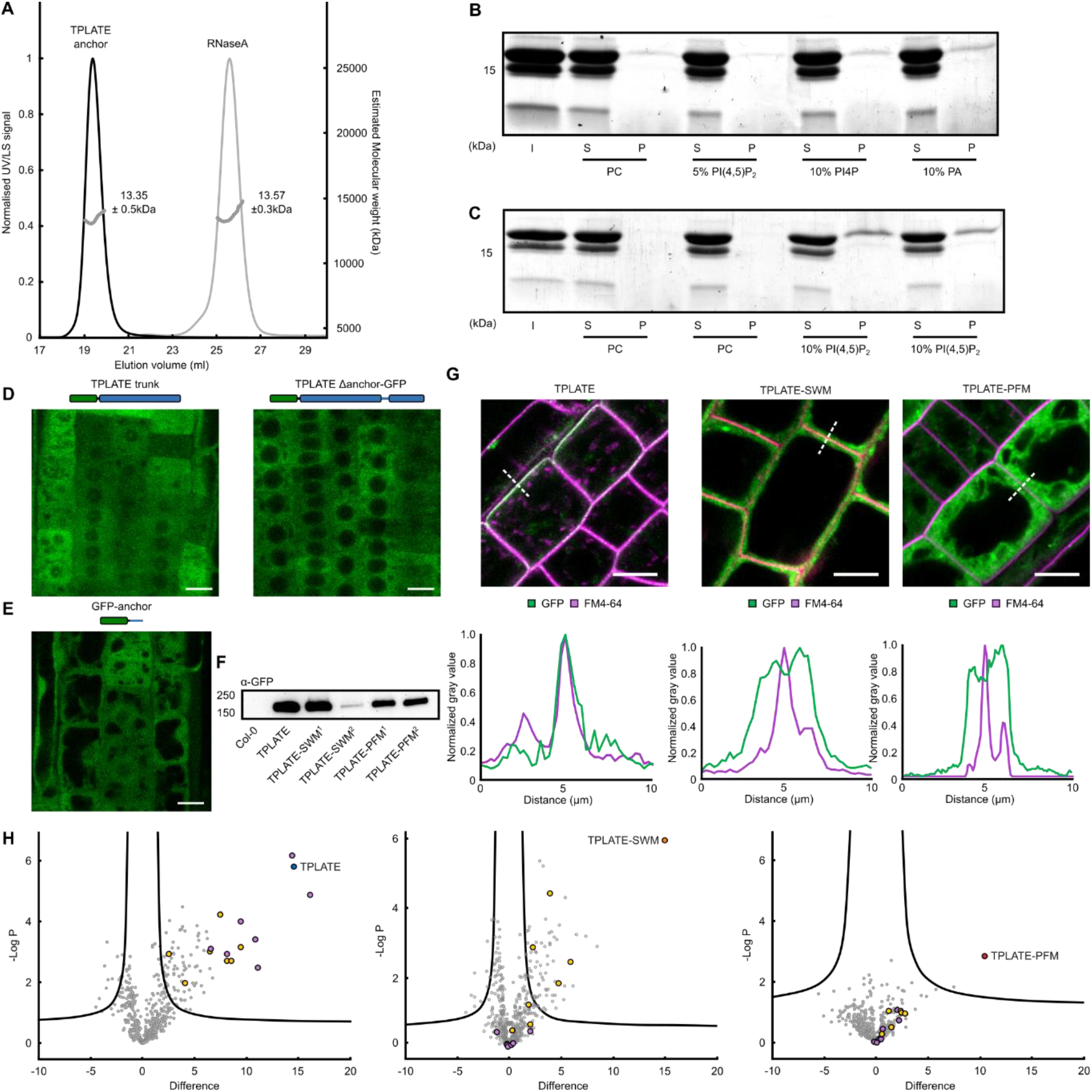
Role of the appendage and anchor domains in targeting and complex assembly. (A) Size exclusion chromatography multi-angle laser light scattering elution profiles (Superdex 75 10/300), showing UV and light scattering signal (LS) of the heterologously expressed and purified TPLATE anchor. RnaseA, which has a similar size, was used as control. The molecular weight distribution (MW) over the main peak is shown as grey dots. (B) Coomassie-blue stained SDS-PAGE analysis of liposome binding comparing the binding of the TPLATE anchor domain between PC/PE and various negatively charged phospholipid containing liposomes. S=soluble fraction, P=pellet. (C) Coomassie-blue stained SDS-PAGE analysis of liposome binding comparing the binding of the TPLATE anchor domain between PC/PE and 10% PI(4,5)P2 containing liposomes. S=soluble fraction, P=pellet. (D) *In vivo* localization of various overexpressed N-terminally fused TPLATE truncations analysed by confocal microscopy. The used constructs are indicated above the image. Absence of PM recruitment of the truncated constructs does not rely on the orientation of tagging as the localization is cytoplasmic, similar to the C-terminally tagged constructs in Figure 5C. (E) The anchor domain does not localize to the plasma membrane *in planta*. (F) Anti-GFP western blot shows the presence of full-length GFP fusions of TPLATE and motif substitution mutants. Two independent lines for each substitution mutant are shown. (G) TPLATE (WT) co-localizes with FM4-64 at the PM, while TPLATE-SWM and TPLATE-PFM mutants display cytoplasmic localization as shown by confocal microscopy. Scale bar is 10 μm. Normalized intensity plots further clarify the presence or absence of the PM localization of TPLATE compared to the TPLATE-SWM and TPLATE-PFM mutants. (H) Volcano plot showing the MS analysis following co-immunoprecipitation of complemented TPLATE as well as TPLATE-SWM and TPLATE-PFM substituted lines compared with Col-0 plants. TPLATE is indicated in blue, TPLATE-SWM in orange, TPLATE-PFM in red and other TPC subunits are indicated in purple. CHCs and DRPs are indicated in yellow. Dots located above the right curve present the significantly detected proteins found when comparing TPLATE or TPLATE substitution mutants with Col-0 using stringent parameters (FDR=0.05 and S0=1).

**Table S1 -.**
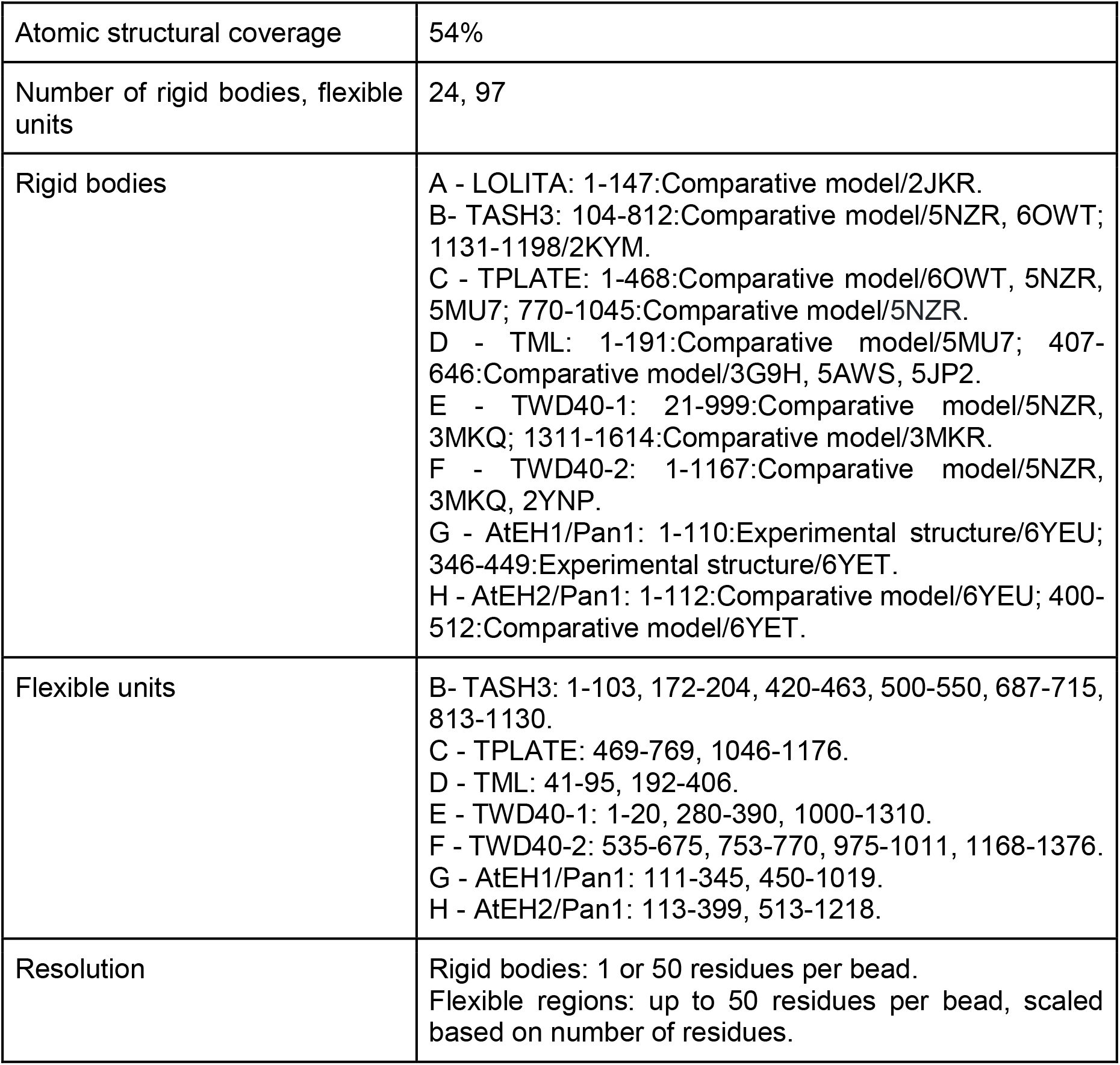
IMP representation of TPC

**Table S2 -.**
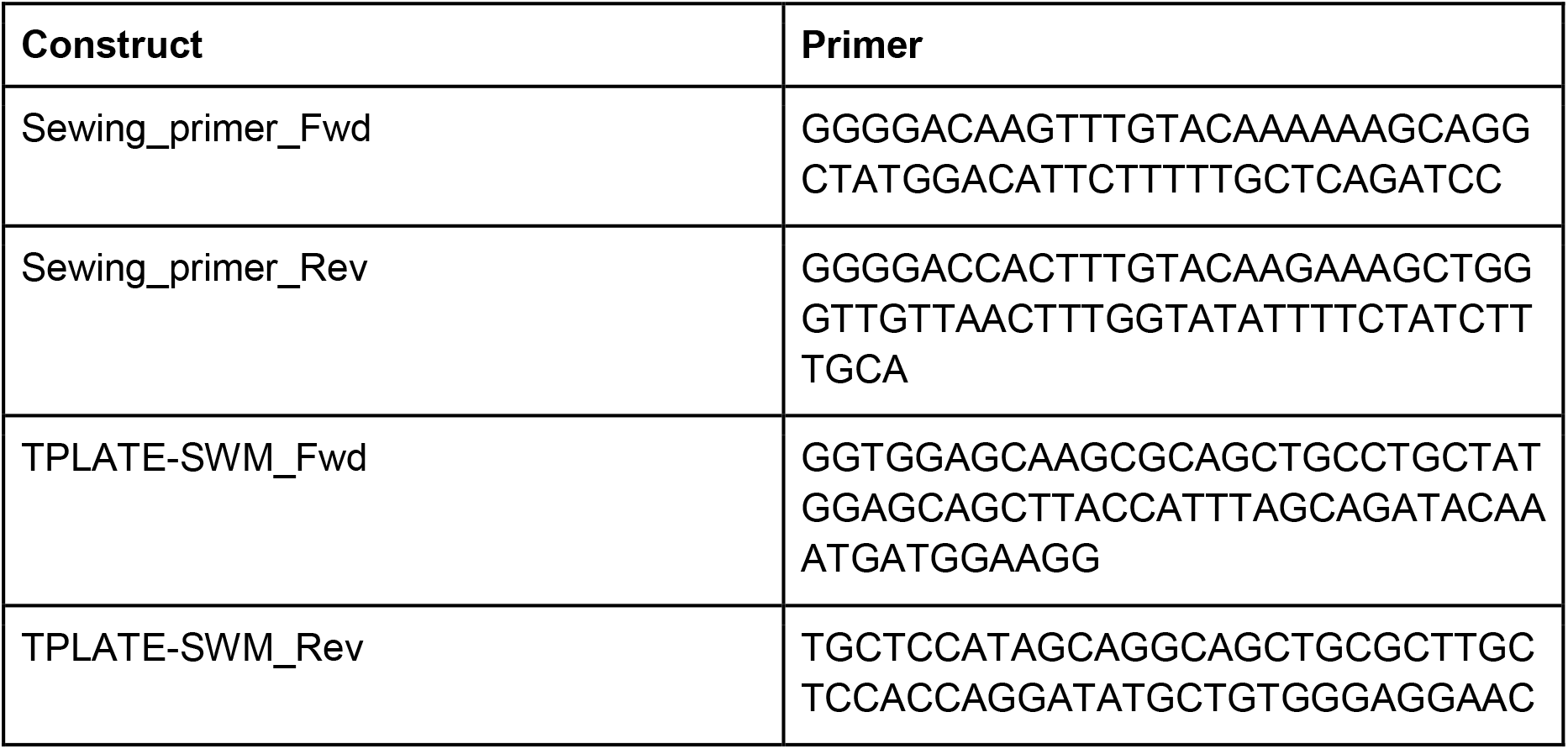
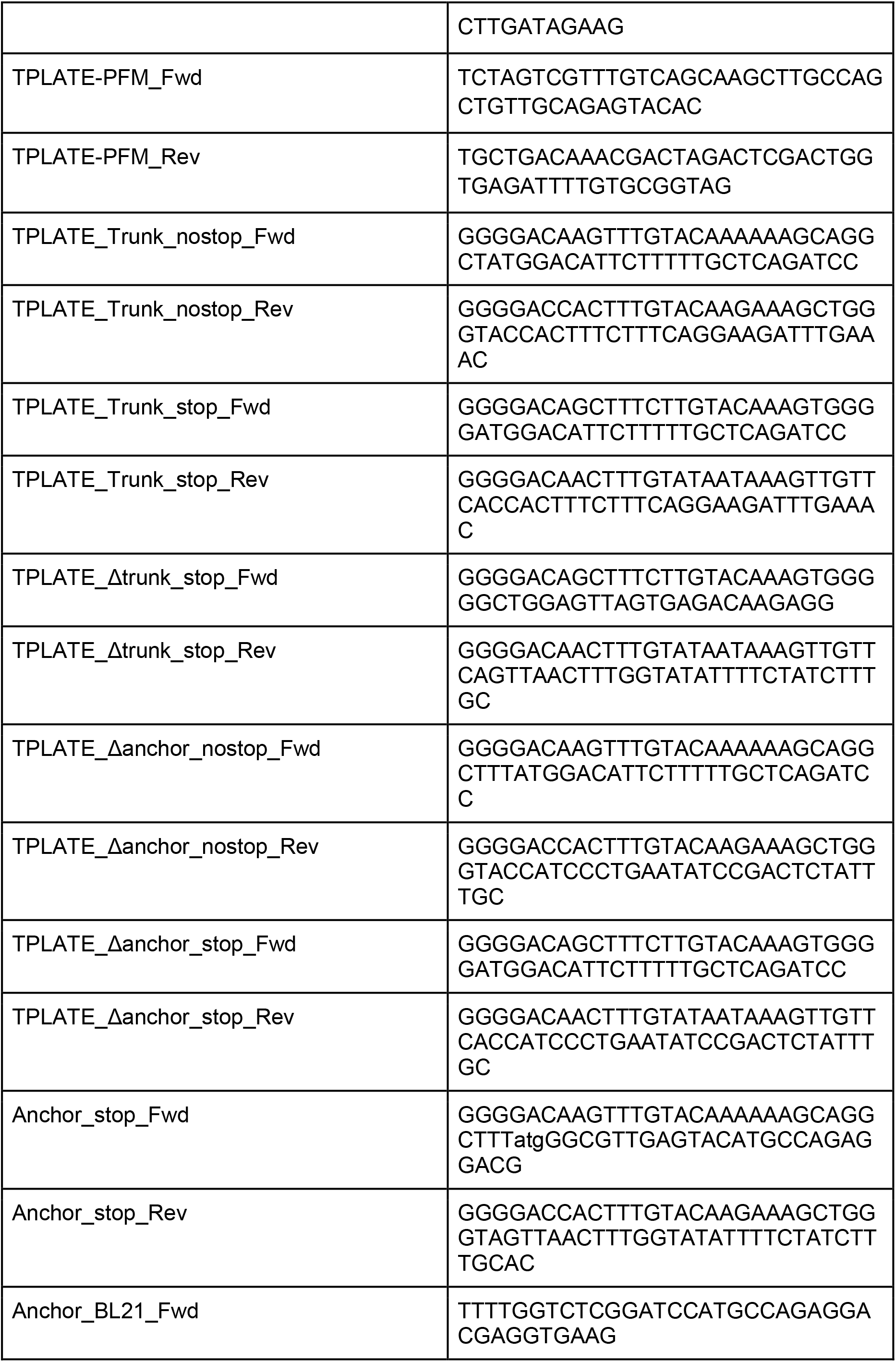
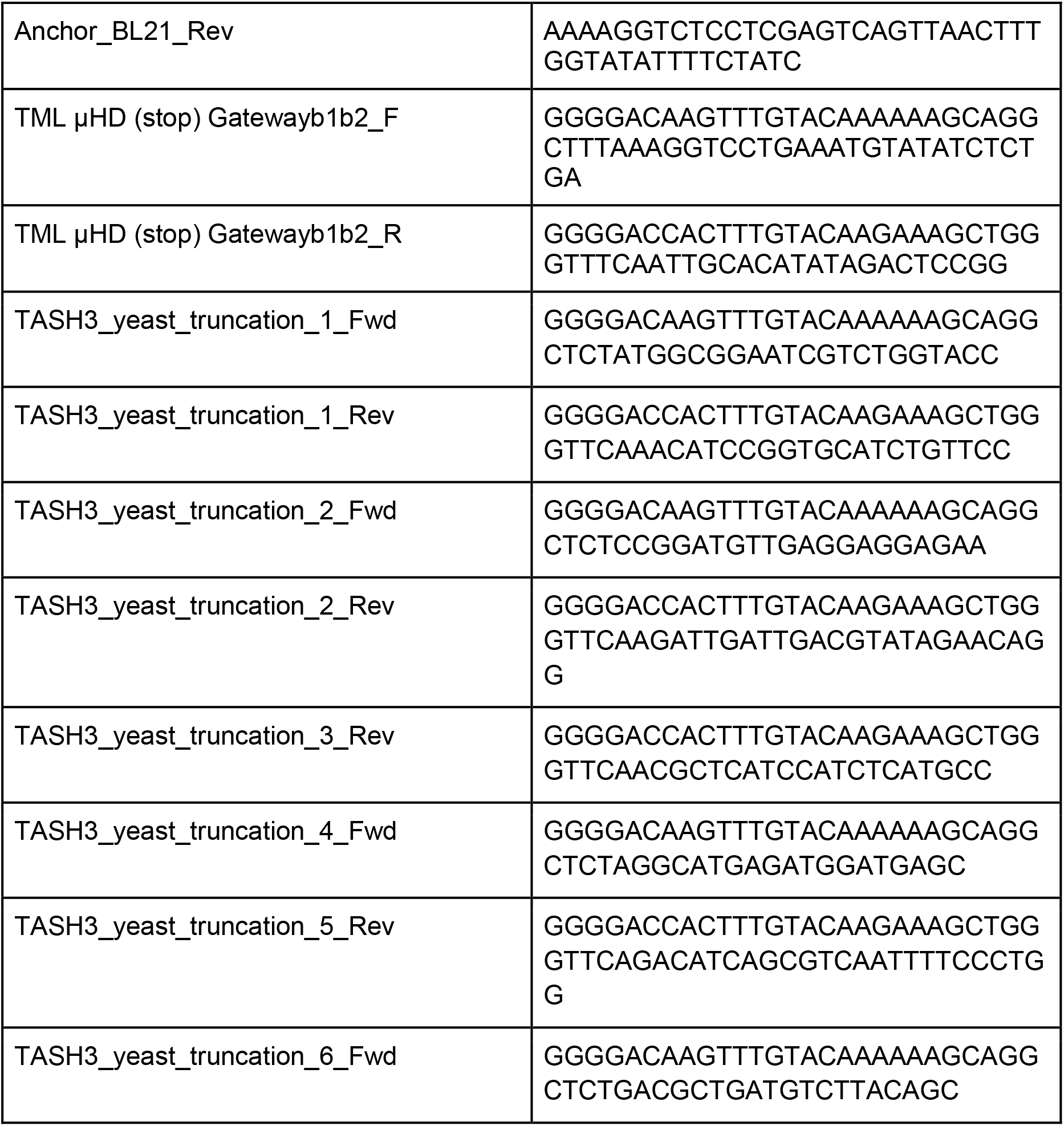
Primers used in the study

**Table S3 -.**
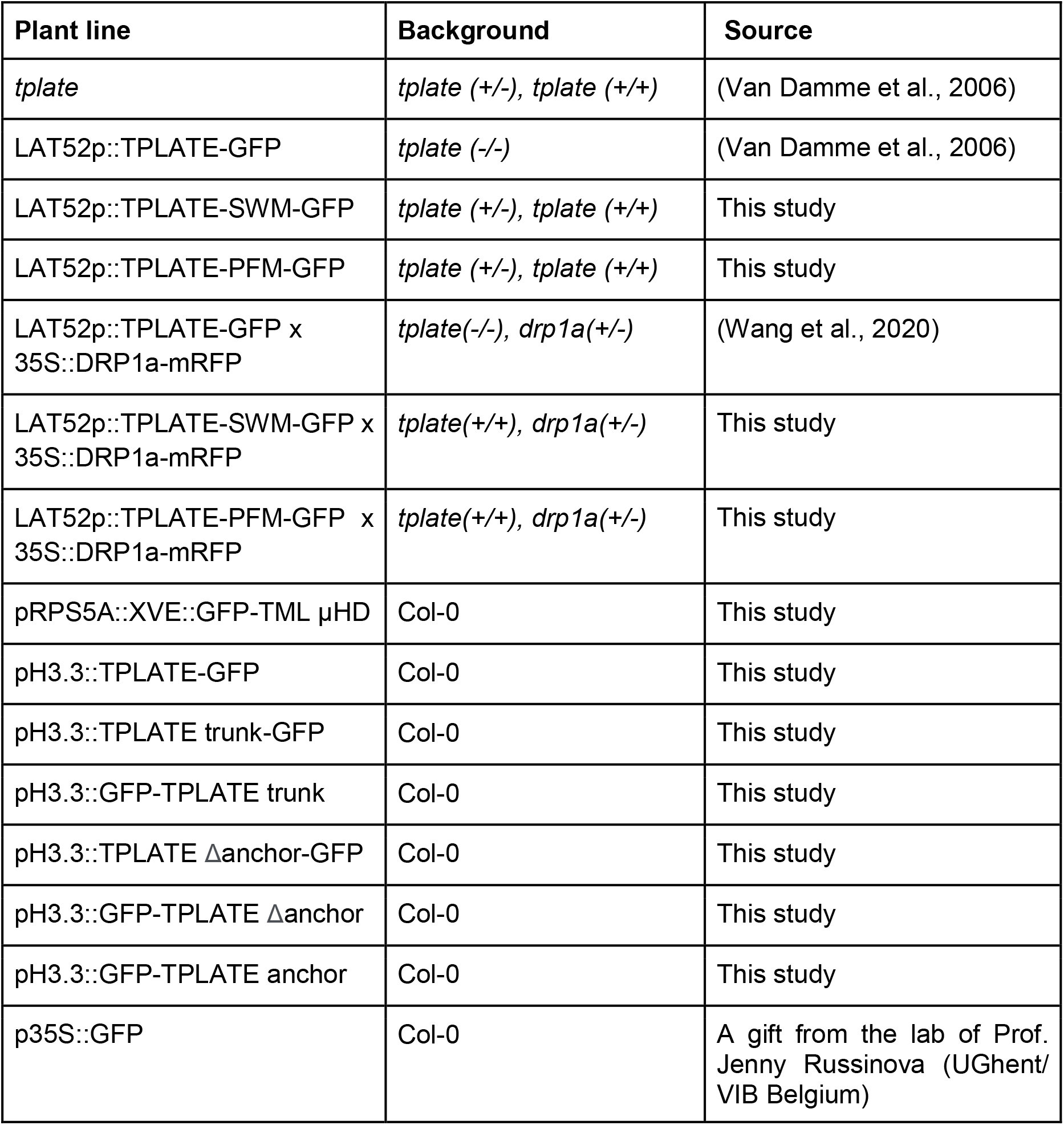
Mutants and transgenic lines used in this study

**Table S4 -.**
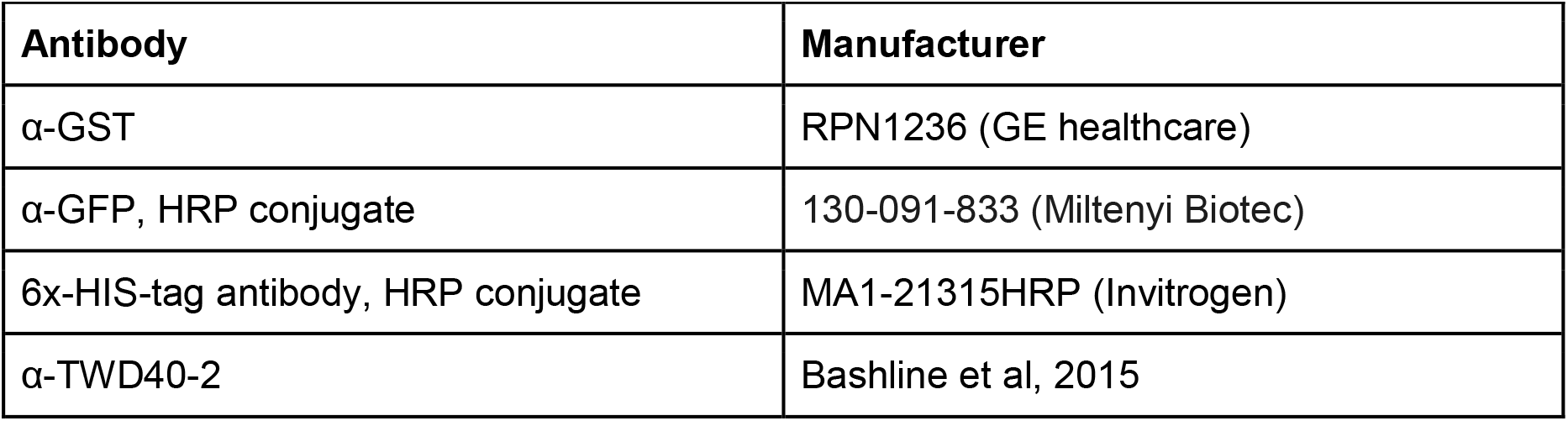
Antibodies used in this study

| Antibody | Manufacturer |
| --- | --- |
| α-GST | RPN1236 (GE healthcare) |
| α-GFP, HRP conjugate | 130-091-833 (Miltenyi Biotec) |
| 6x-HIS-tag antibody, HRP conjugate | MA1-21315HRP (Invitrogen) |
| α-TWD40-2 | Bashline et al, 2015 |

## Materials and methods

### Molecular cloning

Primers used to generate TPLATE appendage substitution fragments (SWM/PFM) are listed in Table S2 and were generated by mutagenesis PCR from the pDONR plasmid containing full-length TPLATE (Damme et al., 2004) by combining sewing and mutation primers after which it was introduced by recombination in pDONR221 via Gateway BP (Invitrogen). All the entry clones were confirmed by sequencing. Expression constructs were obtained by combining the generated entry clones. The TPLATE appendage substituted entry clones were combined with pDONRP4-P1r-Lat52 (Damme et al., 2006), and pDONRP2-P3R-EGFP (Damme et al., 2006) in the pB7m34GW backbone (Karimi et al., 2007) via an LR reaction (Invitrogen). TPLATE truncation constructs as well as full-length were cloned in a similar way but a pH3.3 promoter was used instead. TML μHD was amplified from full-length TML with a stop codon (Gadeyne et al., 2014), cloned into pDONR221 via BP clonase (Invitrogen) and recombined with an RPS5A::XVE promoter and an N-terminal GFP in a pB7m34GW backbone (Karimi et al., 2007) via an LR reaction (Invitrogen) and verified by restriction digest.

Constructs used in the Y2H of the TASH3 truncations were amplified from full-length TASH3 coding sequence by adding gateway sites, the primers are listed in Table S2, and cloned with the help of BP Gateway (Invitrogen) in pDONR221 and verified by sequencing. Entry clones were transformed in a pDEST22 expression vector via an LR gateway reaction (Invitrogen) and checked via restriction digest.

TPLATE Δtrunk was amplified from full-length TPLATE (Damme et al., 2004) after which it was introduced by recombination in pDONR221 via Gateway BP (Invitrogen). Primers are listed in Table S2.

Expression constructs used for KSP were cloned by combining a 35S promoter, entry vectors used in this study as well as FKBP and FRB entry vectors (Winkler et al., 2020). from full length pDONR plasmids. Expression constructs were verified by restriction digest.

### TAP purification and cross-linking MS

Prior to cross-linking, TML and AtEH1/Pan1 subunits were expressed in PSB-D cultures with a C-terminal GS tag and subsequently purified based on an established protocol (Leene et al., 2014). After the purification, the beads were washed with PBS and spinned down for 3min at 500rpm. Fresh BS_3_ cross-linker (A39266, Thermo Fisher) was added and incubated on a rotating wheel for 30min at room temperature. Excess cross-linker was quenched at room temperature for 30min with 50mM NH_4_HCO_3_. For SDS-PAGE analysis, proteins were boiled off from the beads using a mixture of loading dye, PBS and reducing agent. For further analysis using MS, proteins were subsequently reduced with 5mM DTT and acetylated in the dark with 15mM iodoacetamide. Next, the beads were washed with 50mM NH_4_HCO_3_ and incubated overnight at 37 °C with Trypsin/LysC. The supernatant was removed from the beads and desalted with Monospin C18 columns (Agilent Technologies, A57003100) as described in (Leene et al., 2019).

The peptides were re-dissolved in 20 μl loading solvent A (0.1% TFA in water/ACN (98:2, v/v)) of which 10 μl was injected for LC-MS/MS analysis on an Ultimate 3000 RSLCnano system in-line connected to a Q Exactive HF mass spectrometer (Thermo). Trapping was performed at 10 μl/min for 4 min in loading solvent A on a 20 mm trapping column (made in-house, 100 μm internal diameter (I.D.), 5 μm beads, C18 Reprosil-HD, Dr. Maisch, Germany). The peptides were separated on an in-house produced column (75 μm x 400 mm), equipped with a laser pulled electrospray tip using a P-2000 Laser Based Micropipette Puller (Sutter Instruments), packed in-house with ReproSil-Pur basic 1.9 μm silica particles (Dr. Maisch). The column was kept at a constant temperature of 40°C. Peptides eluted using a non-linear gradient reaching 30% MS solvent B (0.1% FA in water/acetonitrile (2:8, v/v)) in 105 min, 56% MS solvent B in 145 min and 97% MS solvent B after 150 min at a constant flow rate of 250 nl/min. This was followed by a 10-minutes wash at 97% MS solvent B and re-equilibration with MS solvent A (0.1% FA in water). The mass spectrometer was operated in data-dependent mode, automatically switching between MS and MS/MS acquisition for the 16 most abundant ion peaks per MS spectrum. Full-scan MS spectra (375-1500 m/z) were acquired at a resolution of 60,000 in the Orbitrap analyzer after accumulation to a target value of 3,000,000. The 16 most intense ions above a threshold value of 13,000 were isolated (isolation window of 1.5 m/z) for fragmentation at a normalized collision energy of 28% after filling the trap at a target value of 100,000 for maximum 80 ms. MS/MS spectra (145-4,085 m/z) were acquired at a resolution of 15,000 in the Orbitrap analyzer. The S-lens RF level was set at 50 and precursor ions with unassigned, single and double charge states were excluded from fragmentation selection.

The raw files were processed with the MaxQuant software (version 1.6.10.43)(Cox and Mann, 2008), and searched with the built-in Andromeda search engine against the Araport11plus database. This is a merged database of the Araport11 protein sequences (http://www.Arabidopsis.org) and sequences of all types of non-Arabidopsis contaminants possibly present in AP-MS experiments. These contaminants include the cRAP protein sequences, a list of proteins commonly found in proteomics experiments, which are present either by accident or by unavoidable contamination of protein samples (The Global Proteome Machine, http://www.thegpm.org/crap/). In addition, commonly used tag sequences and typical contaminants, such as sequences derived from the resins or the proteases used, were added. Parameters Search parameters can be found in Supplemental Dataset 1.

The MaxQuant proteingroups file (Supplemental Dataset 2) was filtered for 2 peptide identifications, and only identified by site, reverse and contaminants were removed. Proteins were ranked by descending iBAQ values, showing that the 8 TPC subunits have the highest iBAQ values, and are thus the most abundant proteins in the samples. Therefore a custom database consisting of the 8 TPC protein sequences was made to use in the pLink2.0 program (Chen et al., 2019). Used parameters can be found in Supplemental Dataset 3. The identified cross-links can be found in Supplemental Dataset 3. The fragmentation spectra of the obtained crosslinks were manually checked and intra-cross-links within 20 amino acids were removed.

### Multiple sequence alignment

Homologues of the TPLATE subunit were taken from published data (Hirst et al., 2014). To identify additional TPLATE subunits homologues, the predicted proteins of each genome were searched using BLASTP (Altschul, S. F. et al. Gapped BLAST and PSI-BLAST: a new generation of protein database search programs. *Nucleic Acids Res*. 25, 3389–3402 (1997)) with Arabidopsis TPLATE as an input sequence. Used databases were GenBank (https://www.ncbi.nlm.nih.gov/genbank/), Joint Genome Institute (https://genome.jgi.doe.gov/portal/), EnsemblPlants (https://plants.ensembl.org/index.html) and Congenie (http://congenie.org/start). Multiple alignment was constructed with the mafft algorithm in the einsi mode (Katoh et al., 2017) and manually normalized on the protein sequence of Arabidopsis TPLATE using the Jalview program (Waterhouse et al., 2009).

### Integrative structure determination of TPC

The *integrative modeling platform* (IMP) package version 2.12 was used (Russel et al., 2012) to generate the structure of TPC. Individual TPC subunits were built based on the experimental structures determined by X-ray crystallography and NMR spectroscopy (Yperman et al., 2020) or comparative models created with MODELLER 9.21 (Šali and Blundell, 1993) based on the the related structures detected by HHPred (Zimmermann et al., 2018) and RaptorX (Källberg et al., 2012). Domain boundaries, secondary structures and disordered regions were predicted using the PSIPRED server (Buchan and Jones, 2019) by DomPRED (Bryson et al., 2007), PSIPRED (Jones, 1999) and DISOPRED (Jones and Cozzetto, 2014).

The domains of TPC subunits were represented by beads of varying sizes, 1 to 50 residues per bead, arranged into either a rigid body or a flexible string of beads (loop regions). Regions without an experimental structure or a comparative model were represented by a flexible string of large beads corresponding to 50 residues each.

For protein-protein docking, a part of a trunk domain of TASH3 or TPLATE was used as a receptor and the entire structure of LOLITA or the longin domain of TML was used as a ligand. Computational rigid-body docking of the respective receptor and ligand was performed using ClusPro2.0 (Kozakov et al., 2017) with default parameters without restraining the interaction site. Docked pairs were then described as a single rigid body for each pair. In total, TPC was represented by 24 rigid bodies and 97 flexible bodies.

119 unique intra and intermolecular BS_3_ cross-links obtained by mass spectrometry were used to construct the scoring function that restrained the distances spanned by the cross-linked residues. The excluded volume restraints were applied to each 10-residue bead. The sequence connectivity restraints were used to enforce proximity between beads representing consecutive sequence segments.

After randomization of position of all the subunits, the Metropolis Monte Carlo algorithm was used to search for structures satisfying input restraints. The sampling produced a total of 1,000,000 models from 50 independent runs, each starting from a different initial conformation of TPC. 4,234 good-scoring models satisfying at least 98% of chemical cross-links were selected for further analysis. To analyze sampling convergence, exhaustiveness and precision, the 4-step protocol (Viswanath et al., 2017) was used. The residue contact frequency map was calculated according to Algret et al. (Algret et al., 2014)

### Yeast two-hybrid (Y2H) assay

Expression vectors were transformed via heat-shock in MaV203. Transforment yeast was grown for two days at 30 °C. Eight colonies were picked up, grown overnight in liquid SD^− Leu/−Trp^ medium and diluted to OD_600_ 0.2 before being plating 10 μl on SD^−Leu/−Trp^ and SD^−Leu/− Trp/-His^ with 50mM 3AT, grown for two days after which the plates were imaged.

### Yeast three-hybrid (Y3H) assay

All TPC subunits were recombined from available gateway entry clones (Gadeyne et al., 2014) in pDEST22 and pDEST32 expression vectors and transformed via heat-shock in both the PJ69-4a and PJ69-4α yeast strains. They were plated out and a single representative colony was picked out and put in culture. A and α strains were mated and cultured in SD^−Leu/− Trp^ and spotted to analyse the Y2H matrix. The liquid cultures were super transformed, via heat-shock, with all TPC subunits (cloned in pAG416GPD) and cultured in SD^−Leu/−Trp/−Ura^. Cultures were grown for two days and were diluted to OD_600_ 0.2 and 10μl was plated on SD^−Leu/−Trp/−Ura^ and SD^−Leu/−Trp/−Ura/-His^ and grown for 3 days at 30 °C after which the plates were imaged.

### Autophagosomal recruitment and Knocksideway in plants (KSP) assay

Tobacco infiltration *Nicotiana benthamiana* plants were grown in a growth room or greenhouse with long-day conditions. Transient expression was performed by leaf infiltration according to (Sparkes et al., 2006). Transiently transformed *N. benthamiana* were imaged two days after infiltration using a PerkinElmer Ultraview spinning-disk system, attached to a Nikon Ti inverted microscope and operated using the Volocity software package. Images were acquired on an ImagEMccd camera (Hamamatsu C9100-13) using frame-sequential imaging with a 60x water immersion objective (NA = 1.20). Specific excitation and emission windows were used; a 488nm laser combined with a single band pass filter (500-550nm) for GFP, 561nm laser excitation combined with a dual band pass filter (500-530nm and 570-625nm) for mCherry and 405nm laser excitation combined with a single band pass filter (454-496nm) for TagBFP2. Z-stacks were acquired in sequential frame mode with a 1 μm interval. Images shown are Z-stack projections. For the KSP assay, An FKBP tagged proteins as well as Mito-FRB and a GFP-tagged subunit were infiltrated. 48 hours after infiltration, *N. benthamiana* leaves were infiltrated with 1 μM rapamycin (Sigma-Aldrich) and imaged in a window between 20-45min. Autophagosomal as well as KSP images were analysed based on the signal in mitochondria versus the cytoplasm (Winkler et al., 2020).

### Coarse-Grained molecular dynamics simulation

The structure of TML μHD was mapped into the MARTINI CG representation using the martinize.py script (Jong et al., 2012; Marrink et al., 2007; Monticelli et al., 2008). The ELNEDYN representation with rc = 0.9 nm and fc = 500 kJ·mol^−1^·nm^−2^ was used to prevent any undesired large conformational changes during CG-MD simulations (Periole et al., 2009). The MARTINI CG model for all lipid molecules used in this study was taken from Ingolfsson et al. (Ingólfsson et al., 2014). Lipid bilayer, in total composed of 600 phospholipid molecules, containing POPC:POPE:POPS:POPA:POPI4P:POPI(4,5)P2 (molecular ratio 37:37:10:10:5:1) was prepared using CharmmGUI Martini Maker (Hsu et al., 2017).

CG-MD simulations were performed in GROMACS v5 (Abraham et al., 2015). GROMACS: High performance molecular simulations through multi-level parallelism from laptops to supercomputers. Softwarex 1–2, 19–25). The bond lengths were constrained to equilibrium lengths using the LINCS algorithm (Hess et al., 1997). Lennard-Jones and electrostatics interactions are cut off at 1.1 nm, with the potentials shifted to zero at the cutoff (Jong et al., 2016). A relative dielectric constant of 15 was used. Simulations were performed using a 20 fs integration time step. The neighbor list was updated every 20 steps using the Verlet neighbor search algorithm. Simulations were run in the NPT ensemble. The system was subject to pressure scaling to 1 bar using Parrinello-Rahman barostat (Parrinello and Rahman, 1981), with temperature scaling to 303 K using the velocity-rescaling method (Bussi et al., 2008) with coupling times of 1.0 and 12.0 ps. Simulations were performed using a 20 fs integration time step. Initially, the protein was placed approximately 3.0 nm away from the membrane. Subsequently, the standard MARTINI water and Na^+^ ions were added to ensure the electroneutrality of the system. The whole system was energy minimized using the steepest descent method up to the maximum of 500 steps, and equilibrated for 10 ns. Production runs were performed for up to 1 μs. The standard GROMACS tools as well as in-house codes were used for the analysis.

### Staining and drug treatment for live-cell imaging

FM4-64 (Invitrogen) was stored at 4 °C in 2 mM stock aliquots in water and protected from light at all times. Whole Arabidopsis seedlings were incubated with ½ MS liquid medium containing 2 μM FM4-64 at room temperature for 15 minutes prior to confocal imaging. Image processing was performed in ImageJ.

PAO treatments were performed for 30 min at room temperature in ½ MS liquid medium containing 30 μM PAO (Sigma-Aldrich).

### Live-cell imaging of Arabidopsis lines

The subcellular localization of TPLATE and TPLATE motif substitutions was addressed by imaging root meristematic epidermal cells of 4 to 5-day-old seedlings on a Zeiss 710 inverted confocal microscope equipped with the ZEN 2009 software package and using a C-Apochromat 40x water Korr M27 objective (NA 1.2). EGFP was visualized with 488 nm laser excitation and 500-550 nm spectral detection and FM4-64 was visualized using 561 nm laser excitation and 650-750 nm spectral detection.

TPLATE truncations were imaged on an Olympus Fluoview 1000 (FV1000) confocal microscope equipped with a Super Apochromat 60x UPLSAPO water immersion objective (NA 1.2). EGFP was visualized with 488 nm laser excitation and 500-600 nm spectral detection.

Dynamic imaging of TPLATE and TPLATE motif substitutions at the PM was performed in etiolated hypocotyl epidermal cells using a Nikon Ti microscope equipped with an Ultraview spinning-disk system and the Volocity software package (PerkinElmer) as described previously (Gadeyne et al., 2014; Wang et al., 2019). Images were acquired with a 100× oil immersion objective (Plan Apo, NA = 1.45). The CherryTemp system (Wang et al., 2020) was used to maintain the temperature of samples constant at 20 °C during imaging.

Seedlings expressing GFP fused proteins were imaged with 488nm excitation light and an emission window between 500 nm and 530 nm in single camera mode, or 500 to 550 nm in dual camera mode. Seedlings expressing mRFP and tagRFP labeled proteins were imaged with 561 nm excitation light and an emission window between 570nm and 625nm in single camera mode or 580 to 630 nm in dual camera mode. Single-marker line movies were acquired with an exposure time of 500 ms/frame for 2 minutes. Dual-colour lines were acquired sequentially (one camera mode) with an exposure time of 500 ms/frame.

### β-estradiol induction

β-Estradiol induction of the pRPS5A::XVE:GFP-μHD line was done by transferring 3-day-old seedlings to medium containing β-Estradiol (Sigma-Aldrich) or solvent (DMSO) as a control. β-Estradiol concentration used was 1 μM.

### Arabidopsis seedling protein extraction

Arabidopsis seedlings were grown for seven days on ½ MS medium under constant light. Seedlings were harvested, flash frozen and grinded in liquid nitrogen. Proteins were extracted in a 1:1 ratio, buffer (ml): seedlings(g), in HB+ buffer, as described before (Van Leene et al., 2007). Protein extracts were incubated for 30 min at 4 °C on a rotating wheel before spinning down twice at 20,000 x g for 20 min. The supernatant was measured using Qubit (Thermofisher) and equal amounts of proteins were loaded for analysis.

### Co-immunoprecipitation assays

Arabidopsis seedling extract, in a 2:1 ratio, buffer (ml): seedlings(g), (see above) was incubated for 2 h with 20 μl pre-equilibrated magnetic GFP-beads (Chromotec, gtma-20). After 2 h the extract was removed and the beads were washed three times with 1 ml of HB+ buffer. Proteins were eluted using a 20:7:3 mixture of buffer: 4x Laemmli sample buffer (Biorad):10x NuPage sample reducing agent (Invitrogen) and incubated for 5 min at 70 °C after which they were loaded on SDS-PAGE gels.

### SDS-PAGE and western blot

Antibodies used in this study are listed in Table S4. Samples were analyzed by loading on 4-20% gradient gels (Biorad), after addition of 4x Laemmli sample buffer (Biorad) and 10x NuPage sample reducing agent (Invitrogen). Gels were transferred to PVDF or Nitrocellulose membranes using the Trans-Blot^®^ Turbo™ system (Biorad). Blots were imaged on a ChemiDoc™ Imaging System (Biorad). Full gels can be found in Supplemental Dataset 7.

### Identification of interacting proteins using IP/MS-MS

Immunoprecipitation experiments were performed for three biological replicates as described previously (Rybel et al., 2013), using 3 g of 4-day old seedlings. Interacting proteins were isolated by applying total protein extracts to α-GFP-coupled magnetic beads (Miltenyi Biotec). Three replicates of TPLATE motif substitution mutants (SWM and PFM) were compared to three replicates of Col-0 and TPLATE-GFP (in *tplate(-/-)*) as controls. Peptides were re-dissolved in 15 μl loading solvent A (0.1% TFA in water/ACN (98:2, v/v)) of which 5 μl was injected for LC-MS/MS analysis on an an Ultimate 3000 RSLC nano LC (Thermo Fisher Scientific, Bremen, Germany) in-line connected to a Q Exactive mass spectrometer (Thermo Fisher Scientific). The peptides were first loaded on a trapping column made in-house, 100 μm internal diameter (I.D.) × 20 mm, 5 μm beads C18 Reprosil-HD, Dr. Maisch, Ammerbuch-Entringen, Germany) and after flushing from the trapping column the peptides were separated on a 50 cm μPAC™ column with C18-endcapped functionality (Pharmafluidics, Belgium) kept at a constant temperature of 50°C. Peptides were eluted by a linear gradient from 99% solvent A’ (0.1% formic acid in water) to 55% solvent B′ (0.1% formic acid in water/acetonitrile, 20/80 (v/v)) in 30 min at a flow rate of 300 nL/min, followed by a 5 min wash reaching 95% solvent B’.

The mass spectrometer was operated in data-dependent, positive ionization mode, automatically switching between MS and MS/MS acquisition for the 5 most abundant peaks in a given MS spectrum. The source voltage was 3.5 kV, and the capillary temperature was 275°C. One MS1 scan (m/z 400-2,000, AGC target 3 × 10^6^ ions, maximum ion injection time 80 ms), acquired at a resolution of 70,000 (at 200 m/z), was followed by up to 5 tandem MS scans (resolution 17,500 at 200 m/z) of the most intense ions fulfilling predefined selection criteria (AGC target 5 × 10^4^ ions, maximum ion injection time 80 ms, isolation window 2 Da, fixed first mass 140 m/z, spectrum data type: centroid, intensity threshold 1.3xE^4^, exclusion of unassigned, 1, 5-8, >8 positively charged precursors, peptide match preferred, exclude isotopes on, dynamic exclusion time 12 s). The HCD collision energy was set to 25% Normalized Collision Energy and the polydimethylcyclosiloxane background ion at 445.120025 Da was used for internal calibration (lock mass).

The raw data was searched with MaxQuant (version 1.6.4.0) using standard parameters (Supplemental Dataset 1). To determine the significantly enriched proteins in bait samples versus control samples, the MaxQuant proteingroups file (Supplemental Dataset 5) was uploaded in Perseus software (Tyanova and Cox, 2018). Reverse, contaminant and only identified by site identifications were removed, samples were grouped by the respective triplicates and filtered for minimal 2 valid values per triplicate. LFQ values were transformed to log2, and missing values were imputated from normal distribution using standard settings in Perseus, width of 0.3 and down shift of 1.8. Next, ANOVA (FDR=0.05, S0=1) was performed on the logged LFQ values, followed by a post-hoc Tukey test (FDR=0.05, Supplemental Dataset 5A). For visualization a hierarchical clustered heatmap was created in Perseus. For visualization as volcano plots (Figure S5-G), t-tests were performed using the logged LFQ values for each bait vs control. The significantly different proteins between bait and control were determined using permutation based FDR. As cut-off, FDR=0.05, S0=1 was applied. Lists of the significantly enriched proteins with each of the baits can be found in Supplemental Datasets 5B, 5C,5D.

### Protein production and purification

TML μHD was cloned into pDEST15 (Gateway). BL21(DE3) cells transformed with the construct were grown at 37 °C until OD_600_ ~0.6 and induced with 0.4 mM IPTG and grown further at 37 °C for 3 hours. Cells were harvested and resuspended in extraction buffer (150mM Tris-HCl pH 7.4, 150 mM NaCl, 1 mM EDTA and 1 mM DTT). The protein was bound on the Glutathione Sepharose (GE Healthcare) matrix and eluted with extraction buffer 1 supplemented with 10 mM glutathione.

TPLATE anchor (1062-1177) was cloned into the in-house generated pET22b-6xHis-TEV. BL21(DE3) cells transformed with the construct were grown at 37 °C until OD_600_ ~0.4-0.6 and induced with 0.4 mM IPTG and grown further at 18 °C overnight. Cells were harvested and resuspended in extraction buffer 2 (20mM HEPES pH 7.4, 150mM NaCl, 1 mM TCEP). The protein was captured on the 5 mL HisTrap HP column (GE/Healthcare) and eluted in extraction buffer containing 150 mM imidazole. The protein was further separated from other impurities by strong anion exchange chromatography using a self-packed Source 15Q column. The 6xHis tag was removed by incubating the protein with TEV protease in a 1:50 (TEV:protease) ratio. After removing TEV by reverse IMAC, the protein was cleaned up using a Superdex 75 Increase 10/300 GL column (GE Healthcare).

### Multi-Angle Laser Light Scattering

Purified His-tagged proteins anchor domain (2.1 mg/ml) was injected onto a Superdex 75 Increase 10/300 GL size exclusion column (GE Healthcare), equilibrated with 20 mM HEPES pH 7.4, 150 mM NaCl and 1 mM TCEP coupled to an online UV-detector (Shimadzu), a mini DAWN TREOS (Wyatt) multi-angle laser light scattering detector and an Optilab T-rEX refractometer (Wyatt) at room temperature. A refractive index increment (dn/dc) value of 0.185 ml/g was used. Band broadening corrections were applied using parameters derived from RNase injected under identical running conditions. Data analysis was carried out using the ASTRA6.1 software.

### Circular Dichroism

The TPLATE anchor domain was buffer exchanged to PBS using size-exclusion chromatography. The protein samples were subsequently spun down for 15 min at 16,200 x g and degassed for 10 min. Far-ultraviolet circular dichroism (CD) spectra were recorded using a Jasco J-715 spectropolarimeter (Tokyo, Japan). CD spectra were measured between 200 and 260 nm, using a scan rate of 50 nm/min, bandwidth of 1.0 nm, and a resolution of 0.5 nm. Six accumulations were taken with samples of TPLATE anchor at 0.2 and 0.4 mg/ml in a 0.1 cm cuvette. The mean residue ellipticity ([θ] in deg · cm^2^ · dmol^−1^) was calculated from the raw CD data by normalizing for the protein concentration and the number of residues using:

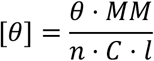

with *MM*, *n*, *C*, and *l*, the molecular weight (Da), the number of amino acids, the protein concentration (mg/ml), and the length of the cuvette (cm) respectively. The secondary structure content was estimated using BeStSel (Micsonai et al., 2015, 2018).

### Lipid-binding experiments

For the liposome binding experiments, either protein-lipid overlay was used according to the manufacturer’s instructions (Echelon Biosciences) or a vesicle co-sedimentation assay was used as described in (Kooijman et al., 2007).

### Statistical analysis

For statistical analysis, the R package in R studio was used. Data were tested for normality and heteroscedasticity after which the multcomp package was used (Herberich et al., 2010).

### Plant Material

Transgenic lines expressing truncation constructs of TPLATE are listed in Table S3. All the plants are in the Col-0 ecotype. The *tplate* heterozygous mutant plants, confirmed by genotyping PCR, were transformed by floral dip with various expression constructs of TPLATE substitution motifs fused to GFP under the control of the pLAT52 promoter, similar to the original complementation approach (Damme et al., 2006). Primary transformants (T1) were selected on ½ MS plates supplemented with 10 mg/L Basta and selected by genotyping PCR to identify transgenic plants containing the *tplate* T-DNA insertion. Genotyping PCR reactions were performed again on T2 transgenic plants expressing TPLATE-SWM or -PFM mutations to identify homozygous *tplate* mutants. Genotyping PCR was performed with genomic DNA extracted from rosette leaves. Genotyping LP and RP primers for *tplate* are described before (Damme et al., 2006), and the primer LBb 1.3 provided by SIGnAL website was used for the T-DNA-specific primer. To obtain dual-marker lines, TPLATE-SWM or -PFM expressing plants were crossed with 35S::DRP1a-mRFP expressing plants (Mravec et al., 2011), respectively. Crossed F1 plants were used for imaging.

### Visualization of protein structures and data

For the visualisation of all protein structures UCSF Chimera (Pettersen et al., 2004) and UCSF ChimeraX (Goddard et al., 2017) were used. Molecular dynamics simulations were visualized with the VMD program (Humphrey et al., 1996). Cross-linking datasets were visualized by xVis (Grimm et al., 2015). All figures were prepared with the Inkscape program (https://inkscape.org/).

### Data availability

All data files related to IMP and CG-MD, and raw MS data are deposited in the Zenodo repository: 10.5281/zenodo.3979550.

## Acknowledgements and Funding

The authors express their gratitude to developers of Linux operating system and open-source programs used in this study, particularly IMP, UCSF Chimera and UCSF Chimera X, Inkscape, ImageJ, GROMACS, VMD, Gimp, Gnumeric, xVIS and Rstudio.

We also like to thank the proteomics core facility of VIB for the help and expertise with running all mass spectrometry experiments.

This research in the D.V.D lab is supported by the European Research Council T-Rex project number 682436 and by the National Science Foundation Flanders (FWO; G009415N). M.P. is supported by the Czech Science Foundation grant GA19-21758S.

## Author contributions

R.P., K.Y. and D.V.D. conceived and planned the experiments.. R.P and K.Y. performed most of the experiments, and wrote the paper together with Ji.Wa. and D.V.D. and with contribution from all co-authors. R.P. and K.Y. performed TAP purification and XL-MS, biochemically characterized TML μHD and performed live-cell imaging. R.P. performed protein-lipid overlay assays, liposome binding assays, integrative structural modelling and molecular dynamics simulations. Ji.Wa. designed and performed complementation assays and did live-cell imaging. D.E. performed MS analysis. Jo.Wi. performed the autophagosomal recruitment assay and knocksideways assay. L.D.V. expressed and characterized TPLATE anchor, and performed the protein-lipid overlay assay. M.V. generated estradiol-inducible lines, performed Y2H and Y3H. P.G. performed co-immunoprecipitation with the TPLATE truncation constructs. E.M. and D.V.D did live-cell imaging. R.P. and M.K. created comparative models. R.M. helped with TPLATE anchor characterization. J.N., E.M. and B.D.R. performed co-immunoprecipitation and MS-analysis of the TPLATE appendage mutants. L.D.V., P.D.B and R.L. performed and interpreted the circular dichroism assay. M.P. helped in designing and performing liposome binding assays. S.N.S., B.D.R., G.D.J., D.V.D. and R.P. were responsible for experimental design, research supervision and finalizing the manuscript text.

